# A central somatotopic map of the fly leg supports spatially targeted grooming

**DOI:** 10.64898/2026.02.27.708590

**Authors:** Leila Elabbady, Grant Chou, Anne Sustar, Andrew Cook, Forrest Collman, John C. Tuthill

## Abstract

Animals continuously monitor their body surfaces to detect and remove debris or parasites. Effective grooming requires that tactile inputs from specific body regions be converted into precisely targeted motor actions, but the neural circuits that support this sensorimotor transformation remain poorly understood. Here, we combine genetic tools and connectomics to elucidate a central somatotopic map of the *Drosophila* leg. We show that the axonal projections of leg touch receptors within the fly’s ventral nerve cord (VNC) are organized along the same cardinal axes as the developing leg. Somatotopically organized bristle axons target a specific class of developmentally-related local interneurons, which imbricate the leg map with overlapping receptive fields of different shapes and sizes. These second-order interneurons target distinct pools of premotor interneurons, which in turn synapse directly onto motor neurons that control leg muscles. Optogenetic activation of second-order interneurons elicits spatially targeted grooming of specific leg regions, consistent with our spatial receptive field predictions based on the connectome. Together, our results suggest that this four-layer circuit processes spatial information from a somatotopic map of the fly leg to guide targeted grooming behavior.

## Introduction

Humans and other animals must constantly monitor the surface of the body to detect and remove unwelcome intrusions. A fly landing on a person’s knee may deflect a hair, which triggers tactile sensory neurons to fire^1^. These signals are transmitted into the spinal cord, where they are transformed across layers of interneurons into patterns of spikes in motor neurons, which move a hand to scratch the leg. Studies in cats and turtles have demonstrated that these animals adapt their scratching movements to reach the site of stimulation^2–4^. This suggests that central circuits are organized to elicit targeted movements in response to activation of specific touch receptors. However, the complexity of vertebrate tactile circuits and the sparseness of previous neuron tracing methods have made it challenging to understand how sensorimotor circuits transform sensory signals into spatially targeted grooming behaviors^5,6^.

A common organizational structure found in early sensory circuits, which may help to simplify such sensorimotor computations, is the somatotopic map^7^. The axons of tactile sensory neurons from neighboring parts of the body often project into neighboring regions of the nervous system forming a map of the body within the nervous system. Within this somatotopic map, the axons from neighboring regions may exhibit similar morphology and connectivity^5,8,9^. In some cases, these sensory maps are preserved in downstream circuits, as in the mammalian somatosensory and visual cortices^10–13^. Understanding the structure of sensory maps is an important prerequisite for deciphering how patterns of sensory neuron activity are transformed into precise motor actions.

In insects, the sense of touch is mediated by tactile bristles distributed across the body^6,14,15^. Each bristle is innervated by a single mechanosensory neuron, which fires action potentials when the bristle is deflected by external forces (Figure 1A). Bristles are extremely sensitive, responding to deflections less than 100 nm^16^. Insects rely on bristles to detect external objects in the environment or debris on their bodies. In *Drosophila*, mechanical or optogenetic stimulation of tactile bristles elicits a range of behaviors including avoidance reflexes and spatially targeted grooming at the site of stimulation^15,17–21^. Some of these spatially targeted behaviors are maintained in headless flies^17,20,22^. This suggests that the fly ventral nerve cord (VNC), the invertebrate analog of the spinal cord, contains the basic circuitry for spatially targeted grooming. Fly grooming is modular and hierarchical: a dirty fly will first clean its eyes and head before proceeding to more posterior body regions like the thorax and abdomen^19,23–26^. Neurons that elicit certain grooming modules (e.g., head, wings, antenna) have been identified^19,27^, but less is known about the neural mechanisms that underlie spatial targeting of grooming movements within a module.

**Figure 1:**
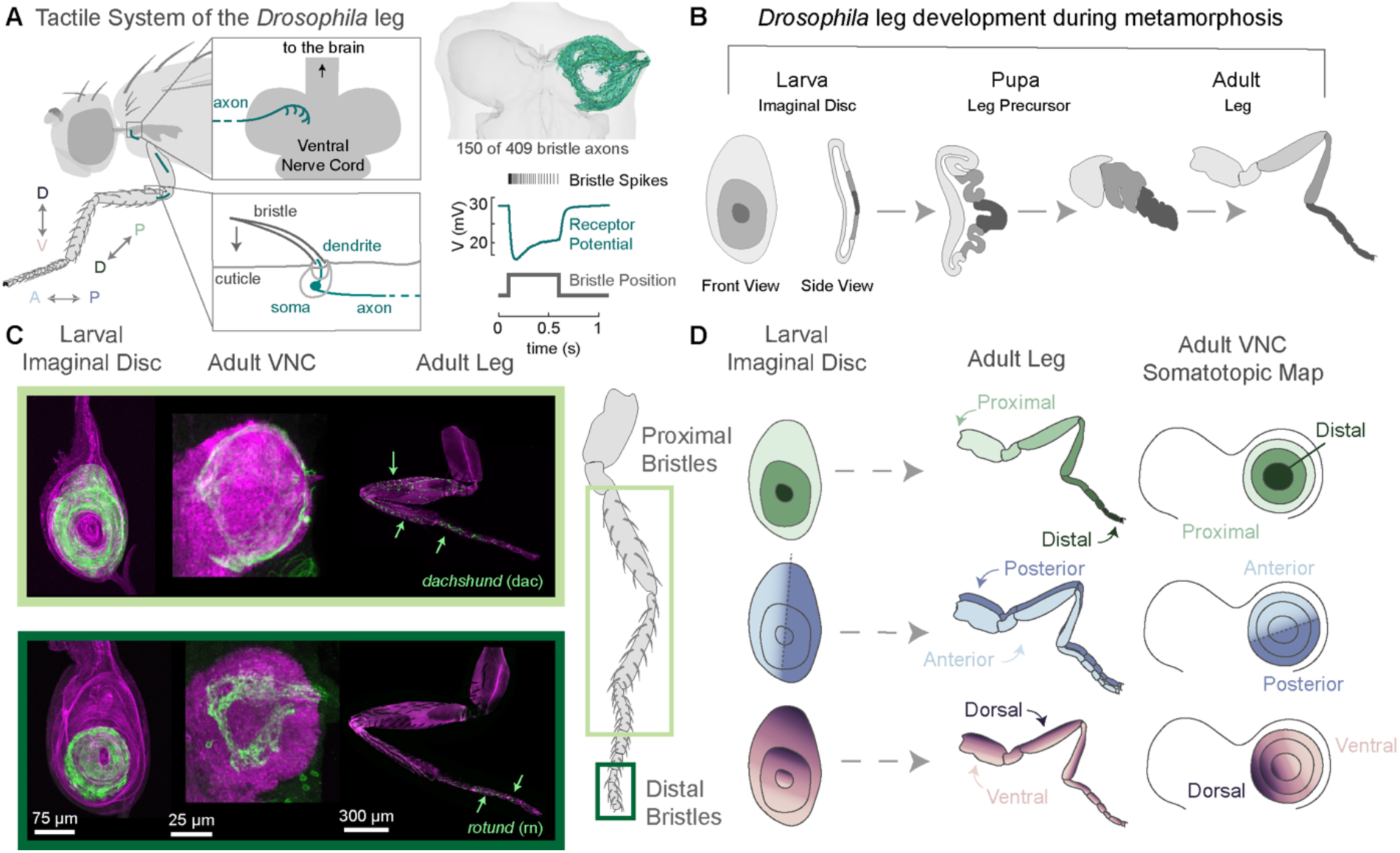
Somatotopy of the leg is maintained in the VNC and recapitulates the larval leg imaginal disc. **A)** A bristle neuron is located at the base of each sensory hair on the leg. The dendrite is stretched upon deflection of the hair (bottom left). Bristle axons project to the ventral nerve cord (VNC) (top left). We reconstructed 409 bristle axons from the front left leg of an adult female fly (top right). Cardinal axes are noted as follows, A: anterior (light blue), P: posterior (dark blue), D: dorsal (dark purple), V: ventral (light purple), P: proximal (light green), D: distal (dark green). **B)** The larval leg imaginal disc develops into the adult leg. **C)** Bristle neurons that express the proximal leg precursor *dachshund* (dac) during development (top). Bristle neurons that express a distal leg precursor *rotund* (rn) during development (bottom). Confocal images show maximum intensity projections of cells in the larval leg imaginal LexAop-mCD8::GFP (green) and an antibody against phalloidin (magenta). Bristle neurons in the leg and VNC were labeled with mcd8::GFP (green) and an antibody against the neuropil marker bruchpilot (magenta). **D)** The somatotopic map of the leg in the VNC recapitulates the somatotopic map of the leg in the larval imaginal disc during development. The proximal to distal axis is mapped along the peripheral to central axis (top). The anterior leg maps to the anterior portion of the VNC leg neuropil and the posterior leg maps onto the posterior leg neuropil (middle). The dorsal leg maps to the area intersecting the anterior to posterior border, while the ventral leg corresponds to axons that remain within either the anterior or posterior region (bottom).

Axons from leg bristles project into the VNC which, like the spinal cord, is organized into neuropil compartments that sense and control specific body parts, including the legs, wings, thorax, and abdomen^15,28–30^. Past work using dye fills of single bristle neurons has revealed that their axons are stereotyped across individuals and suggested the existence of a topographic map within the leg neuropil^5,28,29,31^. However, because each leg has many hundred bristles, the precise organization of the leg map in the VNC remains unknown. Electrophysiological recordings have identified a subset of VNC interneurons that integrate signals from multiple bristle neurons^15^. Yet the circuits that integrate leg bristle signals and transform them into spatially targeted motor commands remain poorly understood.

Advancements in high throughput electron microscopy (EM) and automated image segmentation have resulted in the collection of large volumetric datasets that enable comprehensive cell reconstruction and synapse identification^32–34^. These datasets, referred to as connectomes, enable the study of structural wiring diagrams to predict and understand how circuit architecture facilitates function. Although there exist multiple connectome datasets of the *Drosophila* brain and VNC^35–39^, it remains a challenge to link these connectomes to the fly’s body and peripheral nervous system.

Here, we use a connectome dataset of the *Drosophila* Female Adult Nerve Cord (FANC)^37,40,41^ to investigate how tactile information is mapped in the VNC, from the sensory neurons in the leg through the VNC to the motor neurons that innervate specific leg muscles. We first combined genetic and connectomic tools to elucidate the central somatotopic map of the fly leg. We found that the spatial map of bristle axons in the VNC matches the somatotopic organization of the larval imaginal disc from which the leg develops. We then reconstructed and analyzed how populations of VNC interneurons sample the leg tactile map. Our results suggest a four-layer neural architecture, from leg bristles to motor neurons. Second-order neurons imbricate the leg map into overlapping receptive fields. These second-order neurons target distinct pools of third-order neurons which then target leg motor neurons. Optogenetic activation of second-order interneurons from different regions of the map drove spatially targeted grooming of specific leg regions, consistent with our receptive field predictions from the connectome. Overall, our results elucidate the organization of central circuits in the fly VNC that transform peripheral tactile signals into spatially-targeted behaviors.

## Results

### Leg somatotopy in the VNC recapitulates somatotopy of the developing leg

The front leg of *Drosophila melanogaster* is covered by more than 400 mechanosensory bristles, with the highest density on the more distal leg segments^6^. To understand how tactile information from the leg is mapped in the VNC, we reconstructed 409 bristle axons from the left front leg in a volumetric electron microscopy dataset of a *Drosophila* female adult nerve cord (FANC) (Figure 1A)^37,40,41^. We identified bristle axons based on their morphology and projection patterns into the left front leg neuromere – the region of neuropil corresponding to the left front leg (see Methods). As a population, bristle axons fan out to cover the ventral surface of the VNC; however, each bristle axon innervates a small region within the VNC neuropil. Bristle axons exhibit a range of morphologies (Supplemental Figure 1). While most axons terminate within the same region of the neuropil (e.g., anterior or posterior) there are a subset of axons that branch across the midline in the shape of a hockey stick (Supplemental Figure 1). Across the population, axons with similar morphologies project to similar locations within the VNC neuropil. This structure motivated us to determine the relationship between the location of bristles on the leg and their axonal projections into the VNC.

We developed a genetic strategy to label bristles on specific sections of the leg by restricting the expression of a bristle GAL4 line with transcription factors and signaling molecules that are expressed during development. During metamorphosis, each fly leg develops from an imaginal disc: a cluster of undifferentiated epithelial cells set aside in the embryo and fated to become different parts of the leg (Figure 1B). Graded expression of specific transcription factors and signaling molecules within the imaginal disc regulate the development of the leg along the three cardinal leg axes (anterior/posterior (A/P), dorsal/ventral (D/V), proximal/distal (P/D) (Figure 1A)^42–45^. To take advantage of these spatial patterns, we used a recombinase driven by specific genes to remove a stop cassette and turn on LexAop expression (Supplemental Figure 2A-B, Table 1-2). We applied this method to a suite of genes known to exhibit spatial patterning during development (Table 1), thereby labeling specific bristle cell bodies on the leg and their axons in the VNC. The patterns of spatial expression were sometimes more distributed in the adult VNC than in the larval imaginal disc, so we only analyzed expression from genes that exhibited clear spatial structure in the adult (see methods).

**Table 1:**
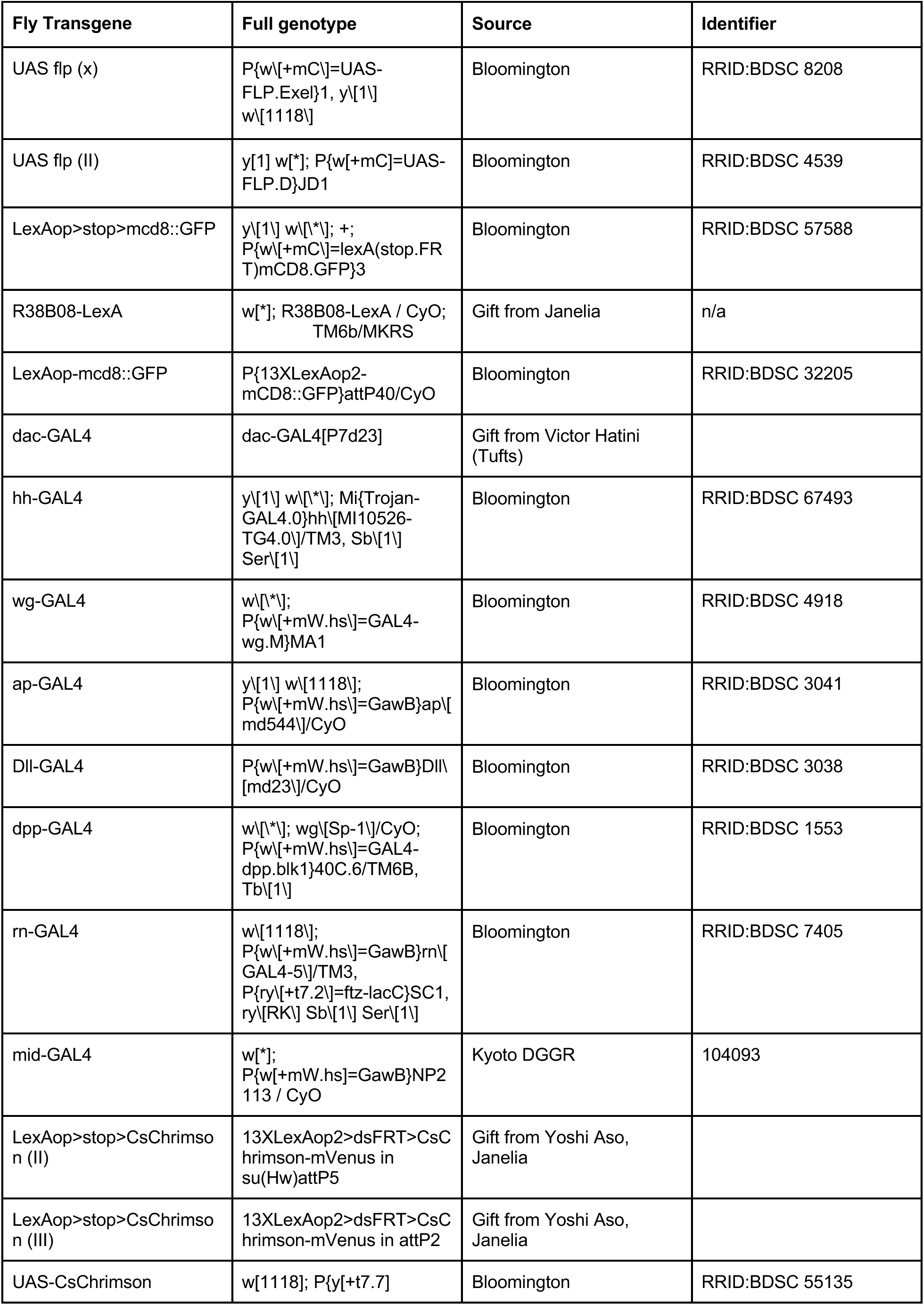
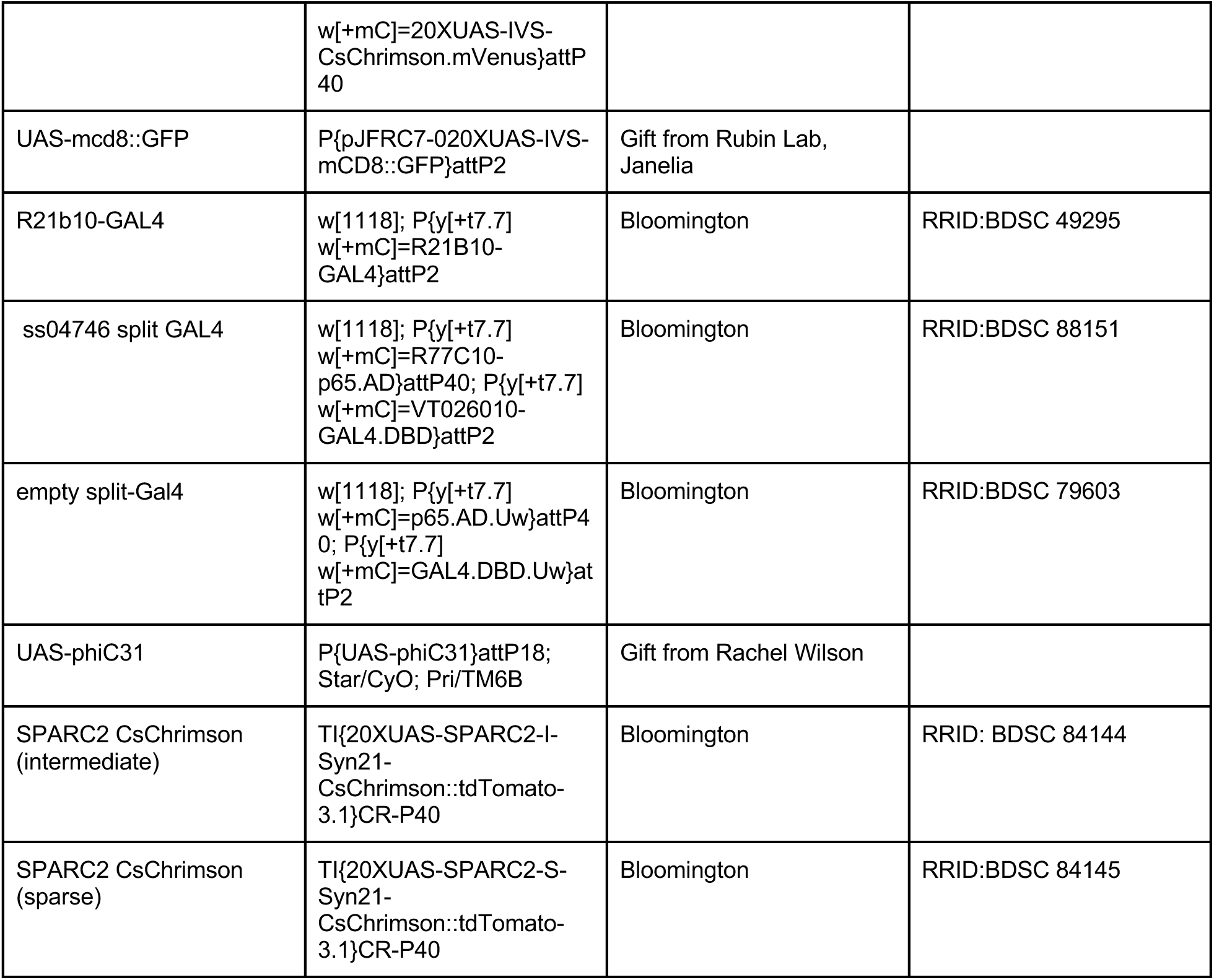
*Drosophila* melanogaster genotypes used for experiments.

**Table 2:** Reagents.

| Reagent | Source | Identifier |
| --- | --- | --- |
| Mouse anti-Bruchpilot antibody | Developmental Studies Hybridoma Bank | RRID:AB_2314866 |
| Chicken GFP polyclonal antibody | Thermofisher PA1-9533 | RRID:AB_1074893 |
| Rabbit DsRed Polyclonal Antibody | Takara Bio 632496 | RRID:AB_10013483 |
| Goat anti-mouse secondary antibody, Alexa Fluor 633 conjugate | Thermofisher A-21050 | RRID:AB_141431 |
| Goat anti-Chicken IgG, Alexa Fluor 488 | Thermofisher A-11039 | RRID:AB_2534096 |
| Goat anti-Rabbit IgG, Alexa Fluor 568 | Thermofisher A-11011 | RRID:AB_143157 |
| Alexa Fluor Phalloidin 647 | Thermofisher A22287 | n/a |
| Vectashield mounting medium | Vector Labs H-1000 | n/a |

Figure 1C shows two examples of transcription factors that label distinct regions of the fly leg and VNC. Bristle neurons that express *dachshund (dac)* during development end up in the proximal leg and project their axons to the outer edges of the VNC neuropil (Figure 1C, Supplemental Figure 2C). Distal leg bristles are labeled by *rotund (rn)* or *apterous (ap)*, and their axons project into the center of the neuromere (Figure 1C, Supplemental Figure 2). Thus, we concluded that the P/D axis of the leg is mapped in concentric rings around the VNC neuropil, with distal bristles at the center and proximal bristles along the outer edges (Figure 1C-D, Supplemental Figure 2C, Table 3).

**Table 3:** Spatial expression of different genes in the adult fly leg.

| Fly Transgene | Gene | Type | Spatial Expression | Spatial Extent on the leg |
| --- | --- | --- | --- | --- |
| hh-GAL4 | Hedgehog (hh) | Signaling molecule | Posterior leg | Femur, tibia, tarsal segments 1-5 |
| mid-Gal4 | Midline (mid) | Transcription factor | Ventral leg | Ventral femur |
| dpp-GAL4 | Decapentaplegic (dpp) | Signaling molecule | Dorsal leg | Dorsal: Coxa, femur, tibia, tarsal segments (1-5) |
| dac-GAL4 | Dachshund (dac) | Transcription factor | Proximal leg | Femur, Tibia, Tarsal segments 1-2 (segments 3 & 4 have one neuron labelled) |
| rn-GAL4 | Rotund (rn) | Transcription factor | Distal leg | Tarsal Segments 1-5 |
| ap-GAL4 | Apterous (ap) | Transcription factor | Very distal leg | Tarsal Segments 4-5 |

During development, the signaling molecule *hedgehog* (hh) contributes to establishing anterior-posterior compartment boundaries in the leg and other body segments^46,47^. Unlike the graded nature of the P/D axis, the distinction between anterior and posterior is defined by a stark compartment boundary. Co-labeling of bristle neurons and hh revealed a similar compartment boundary: bristle cell bodies on the posterior leg, labelled by hh, projected their axons only into the posterior VNC (Supplemental Figure 2, Table 3). For this reason, we defined the A/P axis as a binary map where axons that enter the VNC from posterior portion of the leg neuropil correspond to bristle neurons from the posterior leg and axons in the anterior correspond to neurons in the anterior leg (Figure 1D).

In the leg imaginal disc, cells that express *decapentaplegic* (dpp) develop into the dorsal leg and are located at one end of the imaginal disc at the intersection of the A/P compartment boundary. Co-labeling of bristle neurons that expressed dpp selectively labeled bristle axons that crossed the A/P boundary. These neurons enter the VNC either anteriorly or posteriorly and branch across the midline to the opposing side (Supplemental Figure 1; left column, Supplemental Figure 2C). On the other hand, ventral leg bristle neurons that expressed *midline* (mid) do not cross this border (Supplemental Figure 2C; middle column). In other words, axons that enter the VNC at the anterior edge of the leg nerve terminate anteriorly and vice versa.

In summary, we used spatially restricted labeling of leg bristle neurons along the primary anatomical axes to deduce the somatotopic projections of bristle axons into the front leg VNC neuropil (Figure 1D). Note that for all subsequent analyses, we quantify the A/P axis as a binary proportion. On the other hand, the D/V and P/D axes are more gradual, so we defined the position of each axon along a gradient relative to the population (Figure 1D). The striking similarity between the leg map in the VNC and the imaginal disc (Figure 1D) suggests that the adult leg may develop in coordination with the postembryonic restructuring of the VNC neuropil.

### A predicted axon map recapitulates the distribution of bristles along the leg

Based on the map of the leg we defined through genetic labeling (Figure 1), we developed three mapping rules to predict the peripheral location of bristle axons in the FANC connectome. To mimic the binary compartment boundary along the A/P axis, we defined bristles from the anterior portion of the leg as the axons that arch anteriorly upon entering the VNC from the leg nerves (Supplemental Figure 1; top row), whereas cells located on the posterior leg arch posteriorly in the VNC (Supplemental Figure 1; bottom row). To recapitulate the graded distribution along the P/D axis (Figure 1C-D), we placed a mapping point in the center of the left leg neuropil and calculated the average synaptic distance between each axon and the center mapping point (Figure 2A-B top row, see Methods). Axons that were closer to this mapping point were estimated to be more distal on the leg compared to those further from the mapping point. We used a similar approach for the D/V axis, albeit with a different mapping point to accentuate the relevant axis (see Methods for more details). The spatial predictions for each leg bristle qualitatively matched the patterns observed in genetic labelling experiments (Figure 2B, bottom row). Furthermore, the predicted distribution of bristle neurons along each cardinal axis also recapitulated the expected nonuniform anatomical distribution of bristles along the leg (Figure 2C)^50^. Taken together, these 3D spatial predictions associated each bristle axon in FANC with a relative location along the fly leg and allowed us to trace how spatial information mapped onto the downstream circuitry.

**Figure 2:**
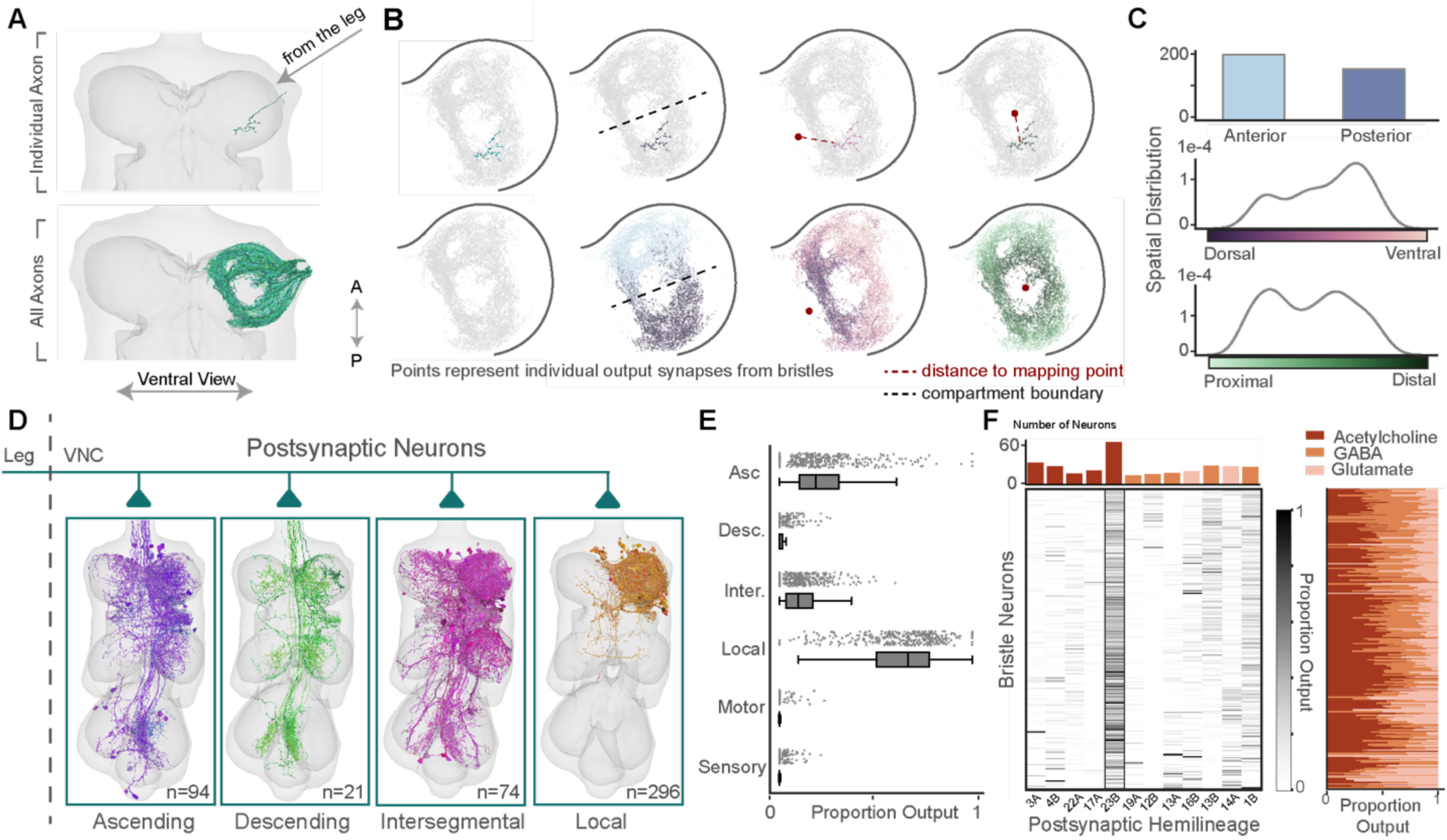
Bristle neurons across the leg preferentially target local 23B neurons in the VNC. **A)** A single bristle axon from the left front leg (top), out of a population of 409 bristle axons reconstructed from the FANC EM dataset, including axons from left front leg nerve, VProN, and DProN^48^ (150/409 axons shown for clarity in the bottom panel). **B)** Output synapses from the single bristle axon shown in panel **A** that arches posteriorly to the A/P compartment boundary, colored by the average synapse distance (d) from the D/V and P/D mapping points respectively (see Methods) (top). Output synapses from all the reconstructed axons colored by their anterior or posterior annotation and the average synapse distance for each individual axon along the D/V and P/D axes (bottom). All synapse points plotted based on their xy coordinates within FANC. **C)** Inferred distributions of bristle neurons on the leg based on the spatial mapping rules. Predicted number of bristle axons on the anterior and posterior leg (top). Predicted spatial density of bristle axons along the D/V axis (middle) and P/D axis (bottom). **D)** Top classes of postsynaptic partners to bristle axons: Ascending n=94, Descending n=21, Intersegmental n=74, and Local n=296. **E)** Proportion output for each bristle axon onto all classes of postsynaptic partners. **F)** Proportion output for each bristle axon (rows, ordered by P/D prediction from top to bottom) onto local VNC neurons from different developmental hemilineages that received on average >=1% of bristle output (columns) (heatmap). Number of unique cells from each hemilineage postsynaptic to bristle neurons (bar chart top). Proportion output for each bristle axon (rows, ordered by P/D prediction from top to bottom) onto postsynaptic partners that release acetylcholine, glutamate, or GABA. Neurotransmitter type assigned based on hemilineage classification for each postsynaptic partner (stacked bar chart, right)^49^. For all box plots, center line, median; box limits, upper and lower quartiles; whiskers, 1.5x interquartile range.

### Bristle axons target local excitatory neurons from the 23B hemilineage

We next used the connectome to analyze the connectivity between bristle axons and downstream neurons in the VNC. Based on automated synapse predictions^37^, each bristle axon makes on average 550 output synapses in the VNC and receives on average 77 input synapses (Supplemental Figure 3). VNC neurons downstream of bristle axons are divided into five broad morphological classes: ascending, descending, intersegmental, local, and motor neurons (Figure 2D, see Methods). On average, local neurons receive the largest proportion of bristle synapses (63%), followed by ascending (22%) and intersegmental neurons (12%) (Figure 2E). Only ∼1% of bristle synapses are onto other sensory neurons. Descending neurons receive less than 2% of bristle synapses, and most bristles make zero synapses onto motor neurons (Figure 2E).

Most neurons in the VNC develop from 33 postembryonic stem cell hemilineages^51^. Cells from the same developmental hemilineage share broad morphological features, typically release the same neurotransmitter^34,49,52^, and may perform similar functions^53^. Using morphological criteria, we classified the developmental hemilineage of each VNC neuron that received input from leg bristles (see Methods). The strongest downstream targets of bristle axons are neurons from hemilineage 23B (Figure 2F). 23B interneurons receive on average 25% of each bristle axon’s synaptic output (Figure 2F, heatmap). Not only are 23B neurons the strongest downstream target, but cells from this hemilineage are the most frequent postsynaptic target of leg bristles (59 cells; Figure 2F, bar chart). Thus, we hypothesized that 23B neurons, as a population, represent a map of the leg and that individual 23B neurons integrate tactile signals from specific regions of this somatotopic map.

### 23B neurons are selective for tactile sensory input

We focused our analysis on 23B interneurons, which release the predominantly excitatory neurotransmitter acetylcholine^49^, because they are the top postsynaptic partner of leg bristles. Of the 59 23B neurons we reconstructed, 56 are local, meaning that their synaptic inputs are restricted to the front left leg neuromere. Two are intersegmental and receive synaptic inputs from multiple leg neuropils and one has an ascending axon that projects to the brain. 23B neurons in the VNC have been well characterized in the literature and are defined by a soma located on the dorsal surface of the VNC, a primary neurite that projects towards the ventral surface with an extensive ipsilateral dendritic arbor that extends through the ventral most surface of the neuropil, and a much smaller contralateral arbor that projects to other segments in the VNC (Figure 3A)^51,53^. In addition to the stereotyped morphology, the primary neurites of the 23B neurons within a segment of the VNC fasciculate together as they enter the neuropil. Regardless of size or morphology, 23B neurons receive on average 40% of their total synaptic input from sensory axons, 85% of which comes from bristle axons (Figure 3B). This suggests that most 23B neurons are specialized for tactile sensing.

**Figure 3:**
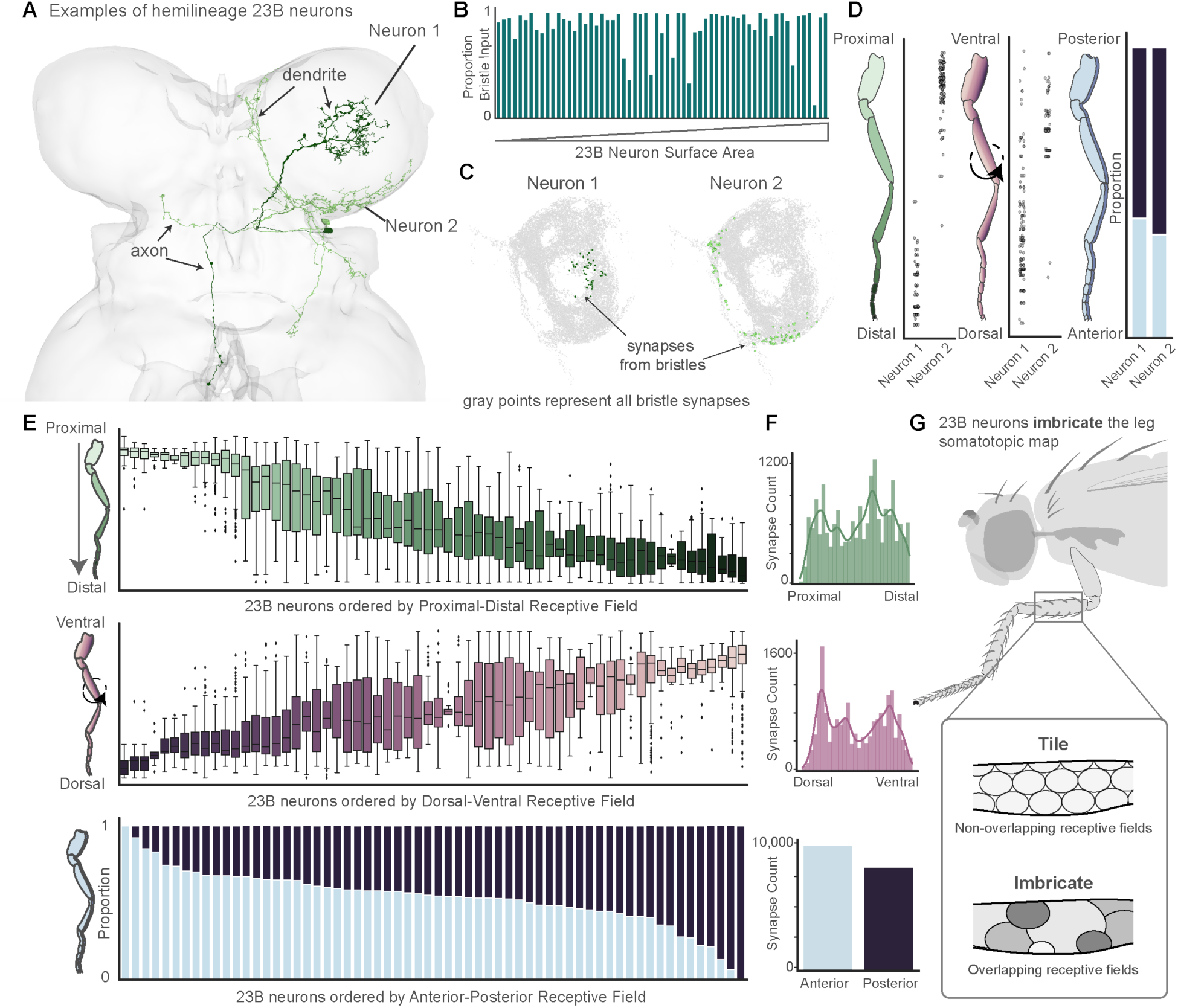
23B neurons imbricate the leg map in distinct overlapping receptive fields. **A)** Two example 23B neurons highlighted in different colors. Dendritic and axonal segments denoted by the arrows. **B)** Proportion of all sensory input from bristle axons for each 23B neuron, bars ordered by surface area. **C)** Input synapses from bristle axons onto Neuron 1 (dark green, n=76) and Neuron 2 (light green, n=136) as compared to all the output synapses from bristle axons (gray). **D)** Receptive field predictions for example Neuron 1 and Neuron 2. Receptive field for the P/D axis (left), D/V axis (middle), and A/P axis (right). **E)** Receptive fields along the P/D axis (top), D/V axis (middle), A/P axis (bottom) for each individual 23B neuron. Individual points represent input synapses from bristle axons and the y axis represents where on the leg each presynaptic bristle axon originates. **F)** Number of bristle input synapses onto all 23B neurons from different areas of the leg along the three spatial axes. For all box plots, center line, median; box limits, upper and lower quartiles; whiskers, 1.5x interquartile range. **G)** 23B neuron receptive fields on the leg imbricate the somatotopic map into overlapping receptive fields, as compared to a non-overlapping tiling pattern.

### 23B neurons imbricate the somatotopic map of the fly leg

Even though all the 23B neurons receive input from leg bristle axons, the dendritic arbors of each 23B neuron within the front left leg neuromere are highly variable (Figure 3A). Based on this diversity, we hypothesized that individual 23B neurons receive input from bristle neurons at different locations on the leg. To quantify this location for each 23B neuron we used the somatotopic mapping approach described above (Figure 2). Each 23B neuron receives input synapses from a selection of bristle axons (Figure 3A, C). Based on our somatotopic mapping, each bristle axon represents a single location on the leg along the three cardinal axes. Therefore, we represented each bristle input synapse onto a 23B neuron by the location on the leg of the presynaptic bristle axon. We refer to the distribution of input synapses along each axis as the receptive field for each 23B neuron (Figure 3C-D). Individual receptive fields varied substantially, as some neurons received input exclusively from proximal or distal bristle axons (Figure 3C,E). Overall, the receptive fields of individual 23B neurons covered the entire somatotopic map of the leg across all three axes (Figure 3D-E). Similar to pebbles on a riverbed, individual 23B neurons *imbricate* the somatotopic leg map by covering the space with overlapping receptive fields of different sizes and shapes (Figure 3G).

### 23B neurons organized by axonal projection patterns

While the dendritic arbors of 23B neurons all project to the ipsilateral leg neuromere (Figure 3), the axons of these cells project to distinct regions of the VNC. Upon closer inspection, we noticed that the axons of 23B neurons that target similar regions in the VNC bundle together within the neuropil. Thus, we reasoned that 23B neurons could be divided into subtypes based on their distinct axonal morphologies. We defined a 23B subtype as a collection of neurons that project their axons to the same region of the VNC: e.g., contralateral T1 (front leg), contralateral T2 (middle leg), or ipsilateral wing. Grouping 23B cells by the axonal projection pattern resulted in 13 subtypes (Figure 4A). Each subtype had between 1-14 neurons. While they were grouped solely by axonal projection, we noticed that 23B neurons within the same subtype had similar dendritic arbors. To quantify this similarity, we represented each 23B neuron by the mean receptive field value in each of the three cardinal axes and calculated the cosine similarity within and between subtypes. We observed that receptive fields were more similar within than across subtypes (Figure 4B-C). The imbrication pattern observed across the entire population of neurons (Figure 3) was maintained at the level of subtypes (Figure 4B). In fact, each 23B subtype occupied a distinct receptive field across the three cardinal axes, suggesting that different subtypes integrate tactile inputs from different parts of the leg (Supplemental Figure 4). Furthermore, we calculated the cosine similarity of postsynaptic partners within and between subtypes and found that the downstream connectivity of 23B neurons was also more similar within subtypes (Figure 4C). This is notable considering that synapses on the axonal projections make up only 17% of 23B output synapses. This means that despite the overlap of dendritic arbors within the left front leg neuromere, 23B neurons from different subtypes target distinct postsynaptic neurons. From these similarities in morphology, receptive field, and postsynaptic targeting, we hypothesized that distinct 23B subtypes function as distinct sensorimotor modules.

**Figure 4:**
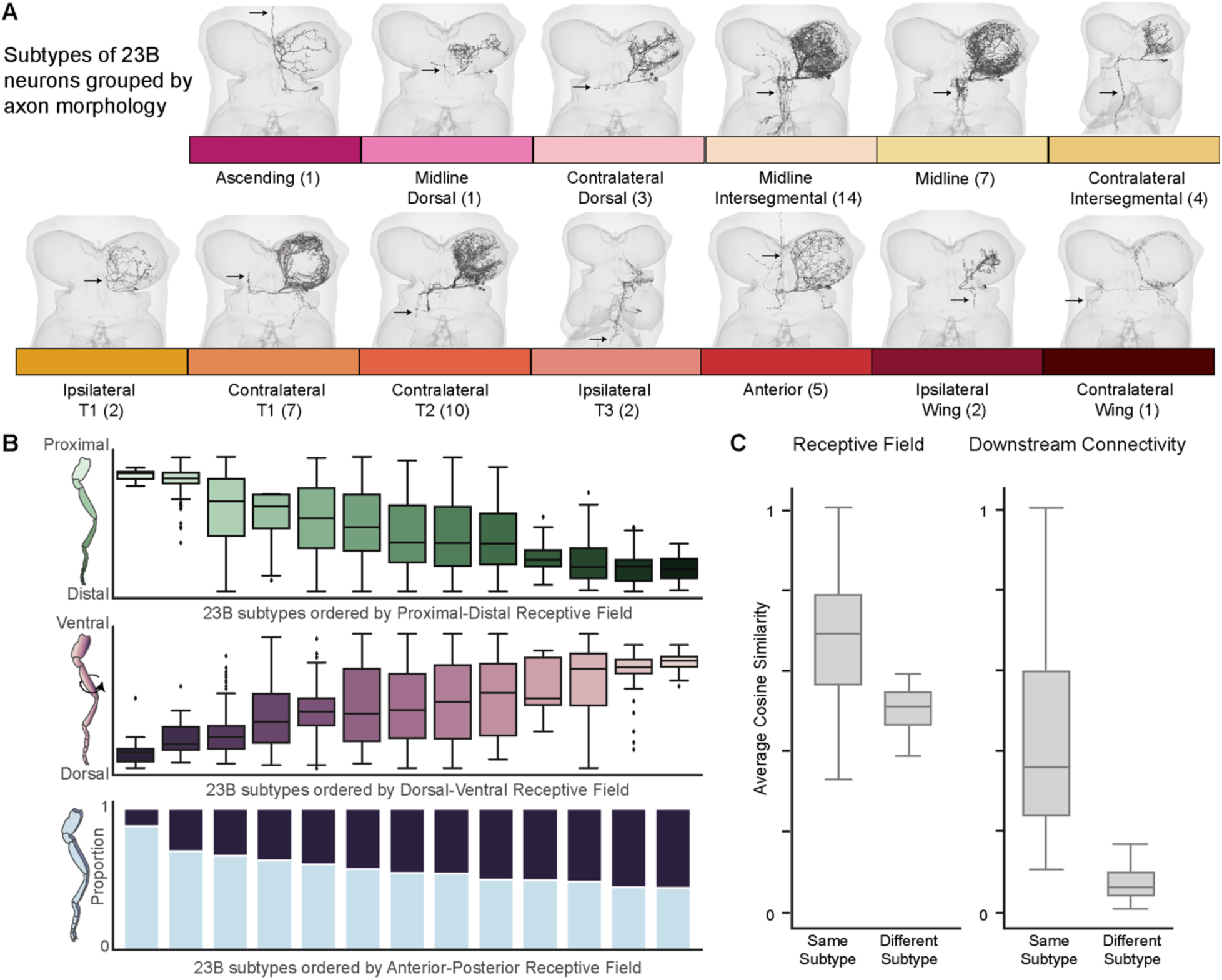
23B subtypes exhibit similar morphology, receptive fields, and downstream connectivity. **A)** 23B neuron morphologies organized and labeled by their axonal projection patterns (arrows indicate axon location). Ascending (1), Midline Dorsal (1), Dorsal (3), Midline Intersegmental (14), Midline (7), Contralateral Intersegmental (4), Ipsilateral T1 (2), Contralateral T1 (7), Contralateral T2 (11), Ipsilateral T3 (3), Anterior (5), Ipsilateral Wing (2), and Contralateral Wing (1). **B)** Average receptive fields of 23B subtypes along the three cardinal axes. 23B subtypes ordered by their receptive fields. **C)** Cosine similarity of individual 23B neurons relative to other 23B neurons within and between subtypes according to mean receptive fields (left) and downstream connectivity (right). For all box plots, center line, median; box limits, upper and lower quartiles; whiskers, 1.5x interquartile range.

### Testing connectome-derived predictions of 23B neuron receptive fields

We used optogenetics to test the behavioral function of 23B subtypes, as defined by their axonal projections (Figure 4). We hypothesized that if 23B neurons are specialized for localizing tactile stimuli, the fly’s behavioral responses to activating these cells would reflect their spatial receptive fields. We identified two genetic driver lines that specifically label distinct 23B subtypes: contralateral T1 (SS04746) and midline intersegmental (R21B10) neurons (Figure 5A, Table 1). We used SPARC^54^ to sparsely label the axons of individual 23B neurons in ∼20 different VNCs for each genetic driver line (Supplemental Figure 6). These sparse labeling experiments confirmed that the two driver lines label different subpopulations of 23B neurons (Figure 5A, Supplemental Figure 6).

**Figure 5:**
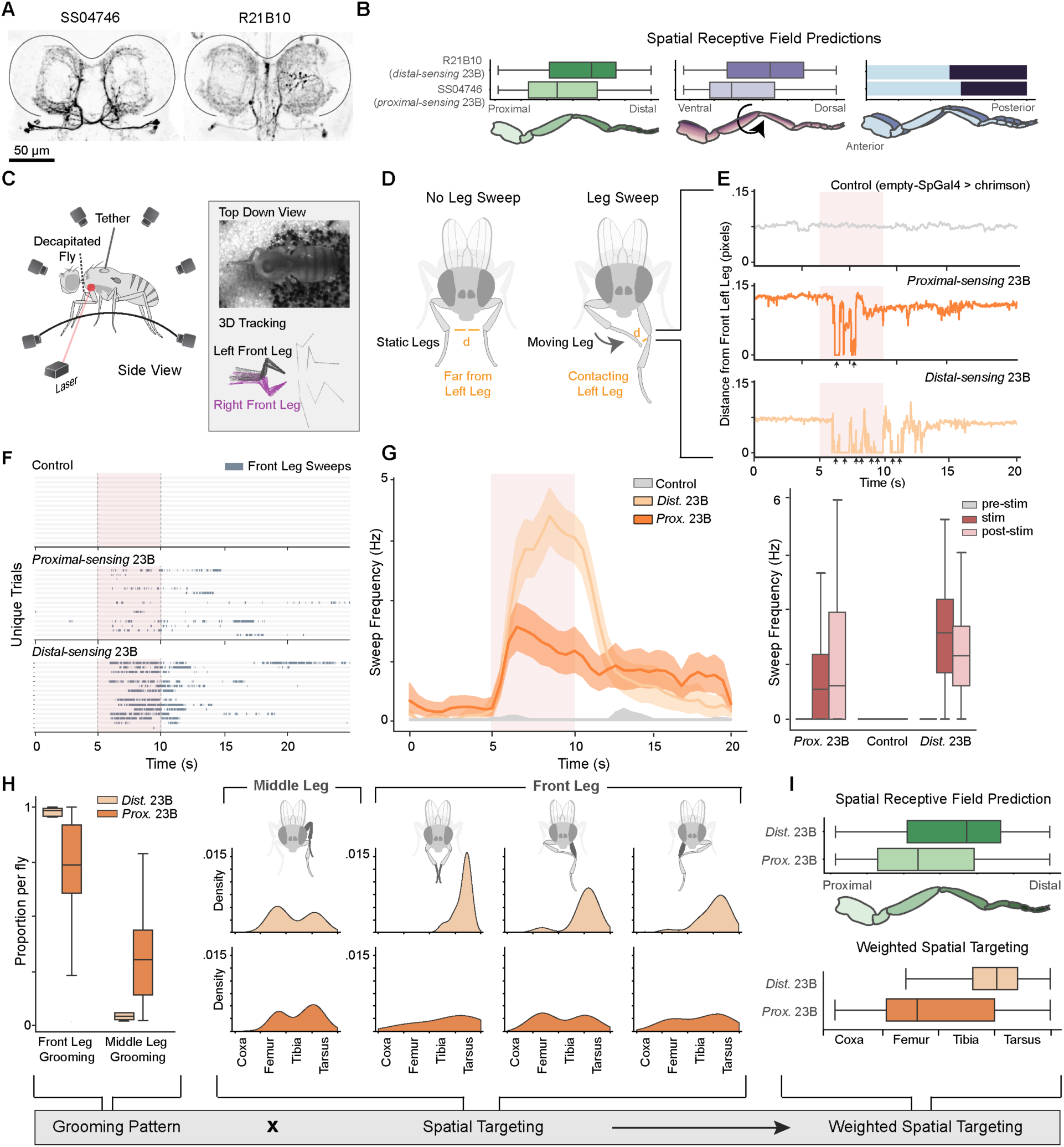
Optogenetic activation of 23B subtypes drives distinct and spatially targeted grooming. **A)** Confocal images show labeling of 23B neurons in the front leg neuropils for two genetic driver lines: SS04746 (left) and R21B10 (right). Neurons labeled with mcd8::GFP (black) (sparsely labeled VNCs in Supplemental Figure 5). **B)** Receptive field predictions for each genetic line across all three cardinal axes (see Methods). Each line is labeled by the predicted receptive field along the proximal-distal axis. **C)** Experimental setup. Headless flies were tethered and positioned on a spherical treadmill. A red laser was focused on the thorax-coxa joint of the left front leg. Behavioral recording and joint tracking was collected from video data from six cameras (inset top) and tracked with DeepLabCut^55^ and Anipose^56^. Bottom inset shows leg movements from one sweep (see Methods). **D)** Individual leg sweeps during grooming were identified as consecutive time points with two legs in close proximity and moving at a minimum velocity (see Methods). **E)** Example trials for empty-SpGal4 flies (control, gray), *proximal-sensing* 23Bs (dark orange), and *distal-sensing* 23Bs (light orange). Distance from the left front leg to the nearest leg over time. Black arrows indicate individually detected sweeps of the left leg. **F)** Leg sweep ethogram with 15 random trials from empty-SpGal4 flies (control, top), *proximal-sensing* 23B flies (middle) and *distal-sensing* 23B flies (bottom). Each row represents an individual trial across time (seconds). Color represents whether the fly was engaged in leg sweeping (blue) or not (gray). **G)** Average sweep frequency (Hz) over time in seconds for control (empty-SpGal4) flies, proximal 23B flies (dark orange), and distal 23B flies (light orange). Distribution of sweep frequency for each line before the stimulus (prestim), during the stimulus (stim), and after the stimulus (poststim). For all box plots, center line, median; box limits, upper and lower quartiles; whiskers, 1.5x interquartile range; outliers not shown. **H)** Proportion of all leg sweeps using front leg grooming or middle leg grooming for each fly (left). Spatial distributions of the first contact point location for a subset of leg sweeps from each grooming pattern (right). **I)** Weighted spatial targeting in response to *proximal-sensing* and *distal-sensing* 23B activation compared to the connectivity-derived receptive fields presented in panel B.

We calculated a connectome-derived receptive field prediction for each genetic driver line. Both SS04746 and R21B10 had six 23B neurons labeled in each neuromere. To predict the cumulative receptive field of these six neurons, we iteratively sampled six neurons from the connectome weighted by the subtype proportions outlined above (Supplemental Figure 5, see Methods). For each sampled subset of 23B neurons, we summed the bristle input from these cells to predict the aggregate receptive field for each driver line (see Methods). From these calculations, we predicted that activation of the 23B neurons in SS04746 would correspond to activation of bristles on the proximal leg and thus elicit proximally targeted grooming. Conversely, the 23B neurons in R21B10 flies received input from distal bristles and thus we hypothesized that activation of these neurons would elicit distal grooming (Figure 5B). Along the D/V axis, we predicted that activating 23B neurons in SS04746 flies would lead to more ventrally targeted grooming compared to 23B neurons in R21B10 flies. Finally, we predicted there would be little to no difference along the A/P axis (Figure 5B). Based on these predictions from the connectome, we refer to the neurons labeled in SS04746 as *proximal-sensing 23B neurons* and the neurons labeled in R21B10 as *distal-sensing 23B neurons*. We hypothesized that driving these different populations of 23B neurons would elicit spatially targeted grooming in line with their respective receptive fields.

For clarity, we reiterate that 23B subtypes are defined by their axonal projections (Figure 4), while the labels *proximal-sensing 23B* and *distal-sensing 23B* refer to the specific 23B neurons labeled by two different genetic driver lines.

### Optogenetic stimulation of 23B neurons drives spatially-targeted grooming

Previous studies have shown that bristle activation in headless flies elicits a range of behaviors including spatially targeted grooming^17,20,22^. This suggests that local VNC circuits are sufficient to support this behavior. To test this more directly, we optogenetically stimulated 23B neurons in decapitated flies while tracking their behavior with 3D pose estimation^56^. To activate 23B neurons in the left front leg neuromere we tethered flies on a spherical treadmill with a red laser targeted at the body-coxa joint of the front left leg (Figure 5C). To quantify spatial targeting of grooming behavior, we identified leg sweeps as consecutive time points where two legs were in contact and moving at a minimum velocity (Figure 5D, individual sweeps noted by the black arrows). Upon activation of distal 23B neurons and proximal 23B neurons, grooming sweep frequency increased (Figure 5E-G, Supplemental Video 1-2). The flies continued to sweep the left leg for several seconds after the stimulus terminated. On the other hand, control flies lacking CsChrimson^57^ expression (empty-SpGal4) did not respond (Supplemental Video 3) (SS04746; 11 flies, 98 trials, R21B10; 8 flies, 73 trials, empty-SpGal4; 10 flies, 80 trials; Figure 5E-G).

### Activation of 23B subtypes elicits different spatially targeted grooming patterns

We observed that the activation of the different 23B subpopulations triggered different grooming patterns. Overall, there were two common grooming patterns in response to 23B activation: grooming the left front leg with the contralateral right front leg (front leg grooming) and grooming the left front leg with the ipsilateral left middle leg (middle leg grooming). *Distal-sensing* 23B activation elicited predominantly front leg grooming (96% front leg, 4% middle leg) while *proximal-sensing* 23B activation elicited both front leg and middle leg grooming (68% front leg, 32% middle leg; Figure 5H).

These results indicate that the fly responds with different behaviors to activation of distinct 23B sub-populations. However, we also sought to determine if the grooming patterns precisely aligned with the predicted receptive field locations of each driver line. For all instances of middle leg grooming, the flies brought the left middle leg forward to rub the stationary left front leg (Figure 5H, left). We observed more variability in front leg grooming so we subdivided these instances into three categories (Figure 5H, see Methods).

To measure the spatial specificity of each grooming pattern, we annotated the first contact position of individual sweeps (see Methods). When we optogenetically activated *proximal-sensing* 23B neurons and the flies produced front leg grooming, the first contacts for each sweep landed on the proximal femur of the targeted leg (Figure 5H bottom). On the other hand, when we activated *distal-sensing* 23B neurons and flies produced front leg grooming, flies targeted the distal portion of the leg, i.e., the tibia and tarsus (Figure 5H, top). *Proximal-sensing* 23B activation also elicited middle leg grooming of the distal femur, tibia, and tarsus, while *distal-sensing* 23B activation triggered middle leg grooming that targeted the middle of the femur. Because flies from the two 23B groups did not use these grooming strategies equally (Figure 5H right), we multiplied the spatial targeting of each pattern by its prevalence to calculate a weighted spatial targeting (Figure 5I, bottom). Overall, we observed that *proximal-sensing* 23B activation elicited grooming of the proximal leg, targeting the middle of the femur (Figure 5I). *Distal-sensing* 23B activation elicited grooming more distally, at the tibia-tarsus joint (Figure 5I). These spatial patterns were consistent with our receptive field predictions based on the connectome (Figure 5I, top).

### 23B neurons do not directly contact leg motor neurons

Activation of both 23B driver lines elicited front leg grooming, however the precise leg movements differed in their spatial targeting (Figure 5H). We therefore wanted to understand how the activation of different 23B subtypes could produce distinct leg movements. In the fly’s front leg, 18 leg muscles are controlled by 71 uniquely identifiable motor neurons^37^. If different 23B neurons produce distinct movements of the same leg, we might expect a difference in their synaptic connectivity onto leg motor neurons. We classified the downstream targets of 23B neurons and the proportion of 23B synapses onto each class type. Other than two cells (both projecting locally to the left front leg neuromere), 23B neurons rarely synapse on leg motor neurons, (1% synaptic output, Figure 6A, Supp Figure 5A). Thus, it is unlikely that 23B neurons directly recruit different leg motor neurons to produce distinct grooming patterns.

**Figure 6:**
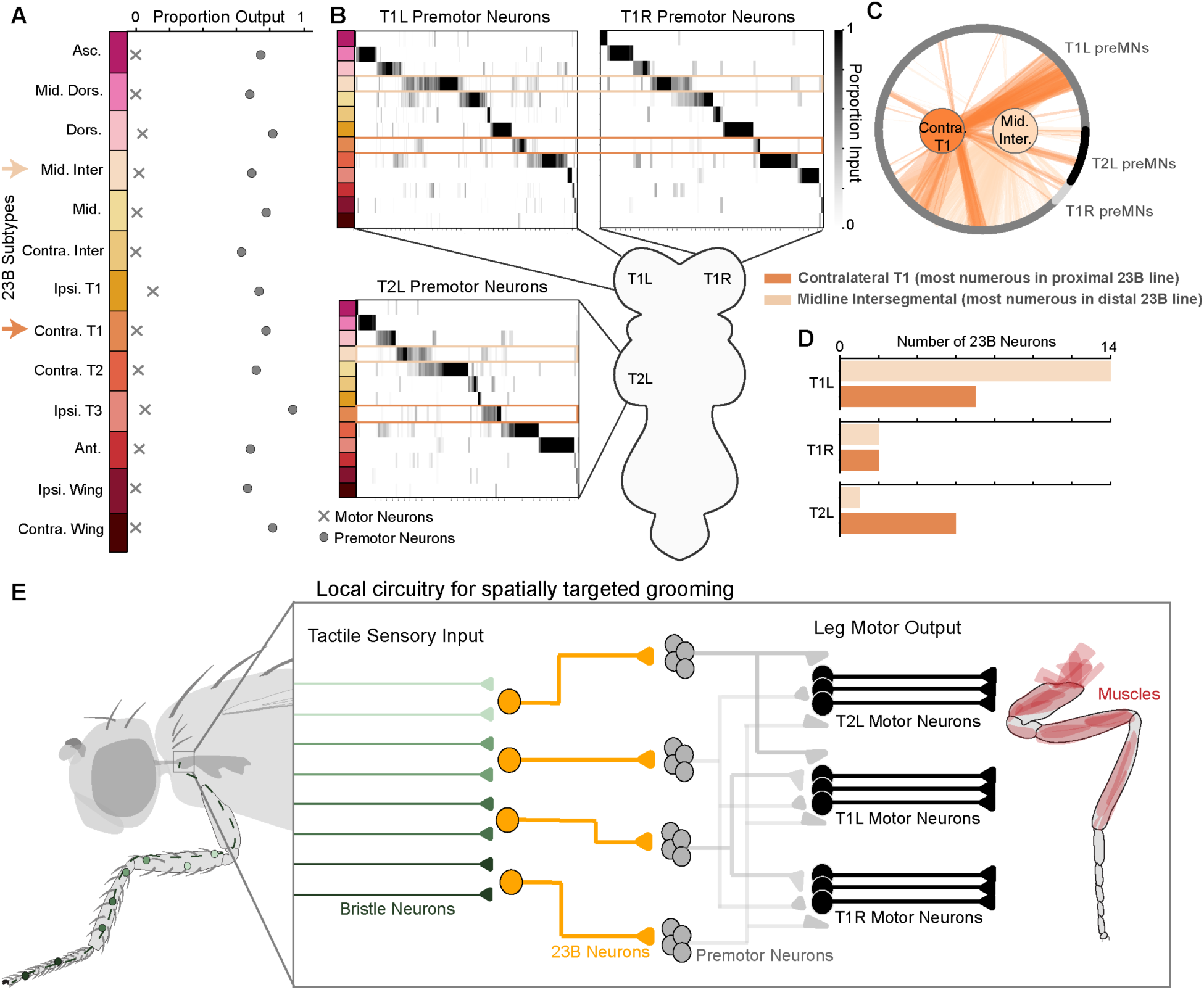
23B subtypes synapse onto distinct leg premotor pools. **A)** Proportion of total synaptic output from 23B neurons onto motor (x) and premotor neurons (o). 23B neurons ordered and colored by subtype. Asc: Ascending, Mid. Dors: Midline Dorsal, Dors: Dorsal, Mid. Inter: Midline Intersegmental, Mid: Midline, Contra. Inter: Contralateral Intersegmental, Ipsi. T1: Ipsilateral T1, Contra. T1: Contralateral T1, Contra. T2: Contralateral T2, Ipsi. T3: Ipsilateral T3, Ant: Anterior, Ipsi. Wing: Ipsilateral Wing, Contra. Wing: Contralateral Wing. **B)** Selectivity of 23B subtypes for left middle leg premotor neurons (T2L), left front leg premotor neurons (T1L), and right front leg premotor neurons (T1R). Colored boxes highlight Midline Intersegmental and Contralateral T1 as the most numerous subtypes in the *distal-sensing* and *proximal-sensing* genetic lines respectively. **C)** Contralateral T1 and Midline Intersegmental subtype connectivity onto T1L, T1R and T2L premotor neurons (preMNs). **D)** Number of 23B neurons from each subtype that contact T1L, T1R, and T2L premotor neurons. **E)** The local four-layer circuit. First-order bristle neurons form a tactile leg map. Second-order 23B neurons imbricate the leg map into overlapping receptive fields and target distinct premotor neuron pools. Premotor neurons recruit leg motor neurons to elicit spatially targeted grooming. Note that recurrence within and between layers is omitted for visual clarity.

### 23B subtypes contact distinct pools of premotor neurons

We next quantified the proportion of 23B target neurons that were premotor. We defined premotor neurons as any neuron that was presynaptic to any motor neuron in the VNC^41^. We further classified each premotor neuron by the motor neurons it targets (e.g., left front leg, right front leg). We found that 75% of 23B synaptic output was onto premotor neurons across the VNC (Figure 6A). In our experiments, we observed that the flies moved the left front leg, right front leg and left middle leg in response to front left leg 23B activation, thus we focused on these three premotor populations for subsequent analyses.

If different subtypes of 23B neurons elicit distinct grooming patterns, then we would expect them to contact distinct populations of premotor neurons. To test this, we measured the proportion of input from each 23B subtype onto individual premotor neurons. We found that across the three leg neuropils (T1L, T1R, T2L), many premotor neurons received input from only one 23B subtype (Figure 6B). While there was some degree of overlap, each 23B subtype synapsed onto a mostly unique set of premotor neurons. This supports the hypothesis that subtypes of 23B neurons recruit distinct motor patterns through distinct premotor populations.

Focusing on the two subtypes of 23B neurons we tested with optogenetics experiments, we observed that these cells contact premotor neurons in three leg neuropils (T1L, T1R, T2L), though the specific populations differ across neuropils (Figure 6B-C). While both 23B subtypes primarily synapse onto left front leg premotor neurons, six of seven contralateral T1 23B neurons make strong connections (12% of their premotor synaptic output) onto left middle leg premotor neurons in T2L (Figure 6D, Supplemental Figure 6B). These results are consistent with our finding that optogenetic stimulation of *proximal-sensing* 23B neurons produced frequent grooming with the middle leg (Figure 5H).

Taken together, we propose that spatially targeted grooming is mediated by a four-layer circuit from tactile sensory neurons to motor neurons (Figure 6E). Tactile sensory neurons target local interneurons that belong to hemilineage 23B. These second-order interneurons imbricate the leg map into overlapping receptive fields and target distinct pools of third-order premotor neurons. Premotor neurons then drive dynamic patterns of leg movement through both direct and indirect excitation and inhibition of leg motor neurons.

## Discussion

### A somatotopic map of the fly leg

In this study, we used genetic labeling to determine that tactile bristles from the fly’s left front leg form a somatotopic map in the VNC. Notably, the spatial patterning of several transcription factors and signaling molecules helped us to infer the somatotopic organization of leg bristle axons in the VNC. The development of the leg is regulated by graded expression of many of the same genes. The P/D and D/V axes of the leg are established by genes like *ap* and *dpp*, while *hh* expression establishes a “compartment boundary” along the A/P axis^46,47^. In the VNC leg neuropil, we found that bristle axons are also organized along a gradient in the P/D and D/V axes, where the projection of each axon is slightly offset relative to its neighbor. Yet the A/P axis is divided by bristle axons that branches either anteriorly or posteriorly, as if separated by a compartment boundary at the center of the left front leg neuromere. In summary, we observe striking similarities between the spatial organization of the leg imaginal disc and the topographic projections of bristle axons in the VNC neuropil.

While the differentiation of the leg imaginal disc occurs well before bristle neurons have developed^58,59^, it is possible that similar molecular factors regulate the temporal differentiation of peripheral sensory neurons and axon guidance into the central nervous system. Recent studies tracing bristle neuron growth and development from the locust antenna suggest that bristle axons enter the nerve tract in order of differentiation^60^. Distal neurons, which differentiate first, enter the nerve tract and are surrounded by more proximal neurons as they grow towards the central nervous system. This results in distal neurons occupying the central region of the tract and proximal neurons concentrically wrapping themselves around the periphery. This topography of the P/D axis is consistent with our findings in the fly VNC and previous work tracing bristle axons from the head^26,61^. While there are many similarities in somatotopic organization of the leg map during development and in the adult VNC, further investigations into the underlying mechanisms and exact timing of sensory axon development in the VNC will be necessary to elucidate how the leg bristle map is established in the fly VNC. Beyond the leg, other precursor structures such as the wing, haltere, and antennal imaginal discs, may also contribute to the creation of somatotopic maps in the adult fly nervous system.

### Grooming behavior

Previous studies mapping the tactile receptive fields of interneurons in the fly and locust proposed that, unlike many other sensory systems, tactile circuits are composed of neural pathways that diverge immediately postsynaptic to sensory axons^8,62^. Our results confirm this hypothesis through the dense reconstruction of the tactile circuit from one leg. We observed that the population of 23B neurons imbricate the leg with distinct yet overlapping receptive fields. After classifying 23B neurons by their axonal projection patterns, we found that neurons of the same subtype contact similar downstream targets and that these different subtypes contact distinct premotor populations across leg neuropils in the VNC. In other words, nearby bristle signals form diverging streams of tactile information that feed into distinct sensorimotor modules. In the spinal cord and the brain, modular motor circuits are found across species and provide a structural scaffold for controlling flexible behavior^63–67^. We propose that distinct 23B subtypes work in concert to activate different populations of premotor neurons that in turn activate motor neurons to elicit targeted grooming responses.

While both the bristle neurons and 23B neurons in the left leg neuropil mainly contact other local neurons (Figure 2E, Supplemental Figure 6A), leg grooming often involves multiple legs. This means that signals from bristles on one leg eventually reach motor neurons in other leg neuropils. For the following reasons, we hypothesize that the intersegmental flow of tactile signals is mediated by 23B neurons: (1) Most 23B neurons project their axons to other VNC neuropils (Figure 4A), (2) The majority of bristle and 23B premotor targets are local neurons (Supplemental Figure 6C), (3) 23B neurons dedicate a larger proportion of their output onto intersegmental premotor neurons than bristle axons (Supplemental Figure 6C). We note that there exist many other intersegmental pathways that could mediate inter-leg coordination of grooming. Although it is not shown in the circuit schematic (Figure 6E), we also note that the post-sensory (i.e., 23B) and premotor layers of the circuit exhibit dense recurrent connectivity. Different approaches will be needed to understand how these circuit motifs support the complex dynamics and contextual gating of grooming and other behaviors involving the legs^41^.

Are grooming circuits for other body parts similarly organized? Previous work in the fly antennal grooming circuit focused on a class of brain interneurons that they refer to as aBN2^19^. Interestingly, the aBN2 cells also develop from hemilineage 23B^61^. aBN2 neurons are strong downstream targets of antennal mechanosensory neurons and, similar to our findings, optogenetic activation of aBN2 neurons increased antennal grooming^19^. These similarities suggest that the structural and functional organization of grooming circuits in the fly may be repeated across body segments. If so, how do these circuits interact when bristles are activated all over the body of a fly? Past work has shown that flies groom their bodies with a stereotyped and hierarchical pattern, starting with the head and proceeding to the legs and abdomen^19,23,24^. Furthermore, several studies have described command-like neurons that elicit grooming of different body segments^19,22,27^. If subtypes of 23B interneurons imbricate each body part, future investigations into the interactions between 23B neurons and these command-like neurons may provide insight into the neural mechanisms that underlie the hierarchical organization and coordination of grooming behavior.

Based on our results and previous studies, we hypothesize that neurons in the 23B hemilineage are involved specifically in the spatial targeting of grooming behavior. Additional evidence for this comes from the fact that activation of 23B neurons in the brain produces targeted grooming of the head and antennae^19^. Second, of all *Drosophila* grooming circuits studied to date, 23B neurons receive the strongest direct input from bristle neurons projecting to both the VNC and brain^61,68^. On the other hand, in addition to grooming, bristle activation can also elicit walking, uncoordinated leg movements, and kicking^17,20^. Here we focused on two 23B subtypes for which we were able to identify specific genetic driver lines, although we predicted the receptive fields for all the 23B neurons (Figure 3). In the future, it will be interesting to explore the range of actions produced by activation of other 23B subtypes, as well as their natural activity patterns during grooming behavior. We note that our approach could be used to define the receptive field of any neuron downstream of bristle axons. In fact, the morphologies of these neurons in FANC suggest that many second-order neurons are subsampling the leg map in the VNC. Characterization of the other interneurons within the tactile circuitry of the VNC will help define the degree to which tactile signals diverge to distinct sensorimotor modules, how this divergence corresponds to the broad range of behaviors elicited from bristle activation, and whether 23B neurons are involved in all spatially targeted behaviors.

### From sensory input to motor output

In our behavioral experiments, we observed several patterns of front leg grooming, suggesting that the spatial location of the tactile stimulus dictates the movement of the leg and patterns of muscle contraction. Our analysis of the sensorimotor circuit suggests that the distinct premotor connectivity of 23B subtypes is important for producing spatially targeted grooming. If bristle neurons can be equated to the pixels of the somatosensory space, we propose that different 23B subtypes sample the leg space to drive the appropriate, spatially-targeted behavioral response. More generally, our work establishes a model circuit within the fly nerve cord to explore how transient sensory stimuli (e.g, touching a leg) produce sustained and dynamic patterns of motor activity.

We have outlined a simplified four-layer circuit (Figure 6E) that includes the major feedforward connectivity of the leg grooming circuitry. However there are three circuit mechanisms that may add complexity and flexibility to the system. The first is that there exist multiple circuits downstream of leg bristle neurons. While 23B neurons are by far the strongest and most numerous downstream partners, there are hundreds of other interneurons that receive direct synaptic input from bristle neurons (Figure 2). Future investigations into how these other interneurons contribute to the range of motor behaviors elicited by bristle neuron activation will elucidate how many parallel circuits are involved in tactile sensation in the fly.

Second, we focused on the excitatory 23B neurons in this study, yet the second strongest target of bristle axons were inhibitory neurons of hemilineage 1B (Figure 2F). Future analyses describing the receptive fields of 1B neurons and their connectivity within the circuit will further our understanding of how inhibitory signals sculpt spatiotemporal processing of tactile signals. Deeper investigation into the balance of excitation and inhibition within the tactile circuit may suggest circuit mechanisms for action selection among the parallel circuits.

Thirdly, while we propose a four-layer circuit as the base, recurrent connectivity is abundant, especially among leg premotor neurons^37,41^. Understanding how tactile stimuli elicit dynamic motor patterns will require recordings of activity dynamics in 23B neurons and downstream cells during behavior. Due to the breadth and variability of premotor populations, experimental perturbations of specific premotor pools and their recurrent connections is currently infeasible. However, dynamic modeling of the connectome has reproduced the functional role of previously characterized cells and revealed the function of uncharacterized circuits^69–72^. As such, *in silico simulations* of the connectome provide an opportunity to directly probe the functional role of specific premotor pools and their respective recurrent connectivity. Future studies that simulate the tactile circuitry could compare how motor neuron outputs change as a function of which premotor pools are activated, elucidate the impact of recurrent connections at each step within the circuit, and measure the influence of initial limb position by manipulating proprioceptive input.

### Limitations

Our connectome results come from one dataset of a female adult nerve cord (FANC). However, the general distribution of bristle axons and the strong downstream connectivity onto 23B neurons is maintained in a connectome dataset from the male adult nerve cord (MANC)^68^. Moreover, we were able to visually identify equivalent 23B subtypes in MANC. This suggests the circuitry is stereotyped across flies and not sexually dimorphic. Similar to conclusions from a comparison of multiple fly brain connectomes^36^, we expect that the overall structure of the bristle sensorimotor circuit is similar across individuals, while the precise connectivity between individual neurons may vary. The consistency between the light-level morphologies described here and the connectome morphologies supports this view, as does the fact that our predictions based on the connectome of one fly were validated in behavioral experiments done on other flies. With the recent availability of connectomes of the full central nervous system^39,73^, future analyses may also elucidate how the connectivity to and from the brain affects grooming dynamics.

## Materials and Methods

### Sample preparation for confocal imaging of imaginal discs

For confocal imaging of imaginal discs (Figure 1, Supplemental Figure 2), we crossed flies carrying the Gal4 driver to flies carrying P{13XLexAop2-mCD8::GFP}attP40. Prothoracic leg imaginal discs were dissected from third instar larvae in PBS, and fixed for 20 minutes in 4% paraformaldehyde in PBS at room temperature. Discs were washed and permeabilized 3x in 0.2% Triton X-100 in PBS (PBST) over 1 hour, then incubated in 1:50 phalloidin for 1 hour at room temperature. The discs were rinsed 3x with PBS over 1 hour, then mounted in VectaShield. We acquired z-stacks on a confocal microscope (Olympus FV1000).

### Sample preparation for confocal imaging of VNCs

For confocal imaging of mcd8::GFP-labeled neurons in the VNCs (Figure 1, Supplemental Figure 2), we dissected the VNC from 2-day old female adults in PBS. We fixed the VNC in a 4% paraformaldehyde PBS solution for 20 min and then rinsed the VNC in PBS three times. We put the VNC in blocking solution (5% normal goat serum in PBST) for 20 min, then incubated it with a solution of primary antibodies (chicken anti-GFP antibody, 1:50; rabbit anti-dsRed 1:500; anti-brp mouse for nc82 neuropil staining, 1:50) in blocking solution for 24 hours at room temperature. At the end of the first incubation, we washed the VNC with PBS with 0.2% Triton-X (PBST) three times over two hours, then incubated the VNC in a solution of secondary antibody (anti-chicken-Alexa 488 1:250; anti-rabbit-Alexa 568 1:250; anti-mouse-Alexa 633 1:250) dissolved in blocking solution for 24 hours at room temperature. Finally, we washed the VNC in PBST three times, once in PBS, and then mounted on a slide with Vectashield (Vector Laboratories). We acquired z-stacks of each VNC on a confocal microscope (Olympus FV1000).

We aligned the morphology of the VNC to a female VNC template in ImageJ with the Computational Morphometry Toolkit plugin (CMTK32; http://nitrc.org/projects/cmtk).

### Sample preparation for confocal imaging of bristles on legs

For confocal imaging of mcd8::GFP-labeled bristles in legs (Figure 1, Supplemental Figure 2), we selected prothoracic legs from 2-day old female adults while the flies were anesthetized with CO2. We immediately fixed the legs in 4% formaldehyde in PBS with 0.2% Triton-X for 20 min and rinsed them in PBS three times over 30 minutes. We mounted the legs in VectaShield and acquired z-stacks on a confocal microscope (Olympus FV1000).

Due to differences in expression between the larva and adult fly, and variability in labeling quality, not all genes that were tested were included in our analysis of the leg map. Only genes that exhibited spatial patterning in the adult VNC were used to define the mapping of the three cardinal axes (Figure 1, Supplemental Figure 2). All genes that were tested are listed in Table 1.

### Bristle neuron reconstruction

409 tactile mechanosensory axons were reconstructed from the front left leg in a connectome dataset of the female adult nerve cord (Figure 2, Supplemental Figure 1)^37,40,41^. Reconstruction, referred to as proofreading, was completed using Neuroglancer, an interactive software for visualizing, editing, and annotating 3D volumetric data. Proofreading entailed two types of edits; splitting off neurites that did not belong to the cell of interest and merging segments of the neuron that were falsely missed by the automated segmentation. All edits and annotations to these neurons are hosted and accessible on the connectome annotation versioning engine (CAVE) platform^74^. 394 of the reconstructed axons entered the VNC through the Leg Nerve, eight from the ventral prothoracic nerve and seven from the dorsal prothoracic nerve. A small number (<20) of axons could not be reconstructed due to irreconcilable segmentation errors.

### Spatial mapping in FANC

To project the spatial axes of the leg map onto the bristle axons in FANC, three mapping rules were applied. The first was that each axon was classified as either anterior or posterior based on whether the axon morphology branched anteriorly or posteriorly upon entering the VNC (Figure 2B). The D/V and P/D axes were quantified along a gradient to reflect the distribution observed from the genetic labeling experiments (Figure 1). For each axis, a mapping point was placed within the neuropil and the distance of every synapse from that point was calculated. To account for spatial outliers, we normalized the distribution of distances along each axis by the 1st and 99th percentile. The relative spatial prediction of each axon was the average synaptic distance from each reference point (Figure 2B).

### Analysis of circuit connectivity

To reduce the presence of weak connections and the likelihood of false positive synapse detections, connections with fewer than three synapses between pairs of neurons were filtered out of all analyses, similar to past work^35,36^. We proofread all downstream targets of the bristle neuron and 23B neuron populations that met this synapse threshold.

We classified each neuron by class (local, intersegmental, ascending, descending, sensory or unknown). We defined local cells as VNC interneurons with inputs limited to the left front leg neuromere, whereas intersegmental cells received input from multiple neuropils. Ascending neurons had a soma in the VNC and projected up through the neck connective. Descending neurons did not have a soma in the VNC and consisted of axons that projected down from the neck connective. We defined sensory cells as afferent axons incoming from the peripheral neurons. Finally, we labeled neuronal fragments that could not be reconnected to the larger arbor as Unknown. Synapses that belonged to an ‘unknown’ object were also filtered out of all analyses (6% of the total connectivity).

We classified all VNC neurons in the tactile circuit by developmental hemilineage. Cells within a hemilineage are born from the same post embryonic stem cell and share morphological features, neurotransmitter expression, and broad functional roles within the VNC^49,52,53^. We assigned hemilineage identity based on soma location, fasciculation bundle into the VNC and dendritic and axonal morphology and projection patterns^51,52,68^. We then inferred neurotransmitter identity from the hemilineage classification based on previously published experiments^49,53^. Less than 1% of neurons could not be classified into a specific hemilineage and were filtered out of any analyses that depended on this labeling (Figure 2F).

### 23B subtype classification

We reconstructed 59 23B neurons downstream of bristle neurons from the left front leg in the FANC connectome. This included 58 from the left front leg neuropil, 3 from the left wing neuropil that extended into the left front leg neuromere. We classified 23B neurons into subtypes based on the axonal projection pattern (Figure 4). For example, 23B neurons in the left front leg neuromere with an axon that projected to the front right leg neuromere were considered Contralateral T1 neurons. 23B neurons that projected to the left wing neuropil were labeled as Ipsilateral Wing neurons and so on (Figure 4A) Axons from neurons of the same subtype bundled together in the VNC. Therefore, in cases where neurons had axons with an ambiguous projection pattern, we classified them based on the axons they bundled with.

### Receptive field calculation

Based on the spatial mapping methods outlined above, we mapped a single location on the leg for each bristle axon and its output synapses (Figure 2B). For each 23B neuron, we selected all the input synapses from bristle axons (Figure 3C). The receptive field along each cardinal axis was represented as the distribution of spatial locations as they were mapped to the presynaptic bristles (Figure 3D-E). If for example a 23B neuron received input from three bristles axons that we had mapped to the ventral proximal area of the leg, the receptive field would be represented by the distribution of input synapses from those three axons. The same method was applied to each 23B neurons (Figure 3E).

### SPARC labeling of 23B neurons

To classify the axonal projection patterns of individual 23B neurons labeled by our two genetic driver lines, we crossed *UAS-PhiC31; ss04746-split-GAL4* or *UAS-PhiC31; R21b10-GAL4* females to males carrying the intermediate or sparse variants of SPARC2 CsChrimson (Supplemental Figure 5). We dissected, fixed, stained, and imaged the VNCs as described above. Neurons were classified by manual inspection of the image stacks based on the morphology and projection pattern of the axon SS04746 (n=21) and R21B10 (n=17). (Supplemental Figure 5)

### Connectome derived spatial targeting prediction

Based on the proportions derived from our sparsely labelled VNCs (Supplemental Figure 5C), we sampled a subset of 23B neurons and summed the bristle input from these cells to predict the aggregate receptive field for that set of neurons. For example, for SS04746, there were six neurons labeled in each neuromere so we sampled six neurons with a sampling rate weighted by the proportion of subtypes present in the SPARC2 experiments (pie chart in Supplemental Figure 5). The aggregate receptive field from this set of six neurons was considered one simulated RF. We then simulated 100 RFs to create the average RF for each genetic driver line.

### Optogenetic experiments

Optogenetic experiments were performed on adult female flies that were raised on 35mM in 95% EtOH ATR for 1-3 days, were 2-5 days old, de-winged, and fixed to a rigid tether (0.1 mm thin tungsten rod) with UV glue (KOA 300). These flies were placed onto a spherical foam ball (weight: 0.13 g; diameter: 9.08mm) suspended by air within a dark arena. A red laser (638 nm; 1200 Hz pulse rate; 30% duty cycle, Laserland) was focused on the thorax-coxa joint of the left front leg (Figure 5C). Optogenetic activation experiments were conducted on flies in which different subtypes of 23B neurons expressed CsChrimson, as well as flies with an empty-SpGal4 (**Table 1**)(SS04746; 11 flies, 98 trials, R21B10; 8 flies, 73 trials, empty-SpGal4; 10 flies, 80 trials). Trials were 20 seconds in duration and consisted of five seconds prestimulus, five second with the laser flickering on/off at 5Hz, and 10 second post stimulus (Figure 5E). During each trial, the behavior each fly was recorded with 6 high-speed cameras (300 fps; Basler acA800-510 µm; Basler AG) and the movement of the ball was recorded at 30 fps with a camera (FMVU-03MTM-CS) and processed using FicTrac^75^. The 3D positions of each leg joint were determined by using DeepLabCut^55^ and Anipose^56^ (Figure 5C-D). Kinematic analyses were performed in a custom Python script.

### Leg sweep detection

We used the 3D joint positions to detect contacts between legs (Figure 5C-D). The automated tracking detected the following joints for each leg of the fly: body-coxa, coxa-femur, femur-tibia, tibia-tarsus, and the tarsus tip. We interpolated vectors between the joints of individual legs to represent the legs in 3D space. We defined contacts as individual frames where two legs were in close proximity to one another. The distance threshold we used to classify contacts varied between flies to account for diurnal variability in camera calibration settings, however they all ranged between 0.13-0.17 pixel distance. We defined leg sweeps as consecutive frames with a contact detection between the same two legs. At least one of the legs had to be moving at a minimum velocity of 2 mm per second to be considered a valid leg sweep (Figure 5D-E). We added the velocity condition to exclude moments when the fly idly stood with two legs in contact. Finally, to account for noise from the binary contact detection, we merged individual sweeps that were separated by three or less frames (Figure 5D-E).

### Spatial targeting and contact point annotation

To define the spatial targeting of each grooming pattern we needed the exact contact point location between legs. Since we tracked joint positions and not entire leg segments, we annotated the contact points for a subset of frames that could then be measured relative to our interpolated legs. To do this we defined the first point of contact as the first frame of each individual leg sweep. We then divided first contacts by grooming pattern based on the legs involved; sweeps between the left front leg and the middle front leg were considered middle leg grooming, sweeps between the two front legs were considered front leg grooming (Figure 5H). We sampled first contact frames for each grooming pattern across the two populations of experimental flies: middle leg grooming SS04746 (53), middle leg grooming R21B10 (42), front leg grooming SS04746 (32), front leg grooming R21B10 (67). All frames were annotated by a person blind to genotype using the point annotation software Anivia. We annotated the contact point location across all six camera views for each frame. Due to the variability in front leg grooming we also annotated the category of front leg grooming. We defined Category 1 as both front legs towards the midline. Category 2 was when flies brought the right leg over to the left side and contacted an extended left leg. Category 3 when flies brought the left front leg over to the right and contacted an extended right leg (Figure 5H).

To compare contact point locations relative to the leg in 3D space, we triangulated the annotated contact points into the same space. This was done by importing the calibration settings for each respective trial and running the tracking process described above. To determine the spatial location of the contact we measured the closest point on the interpolated legs to the annotation point. We defined the spatial targeting profile as the distribution of leg locations contacted for each grooming pattern (Figure 5H).

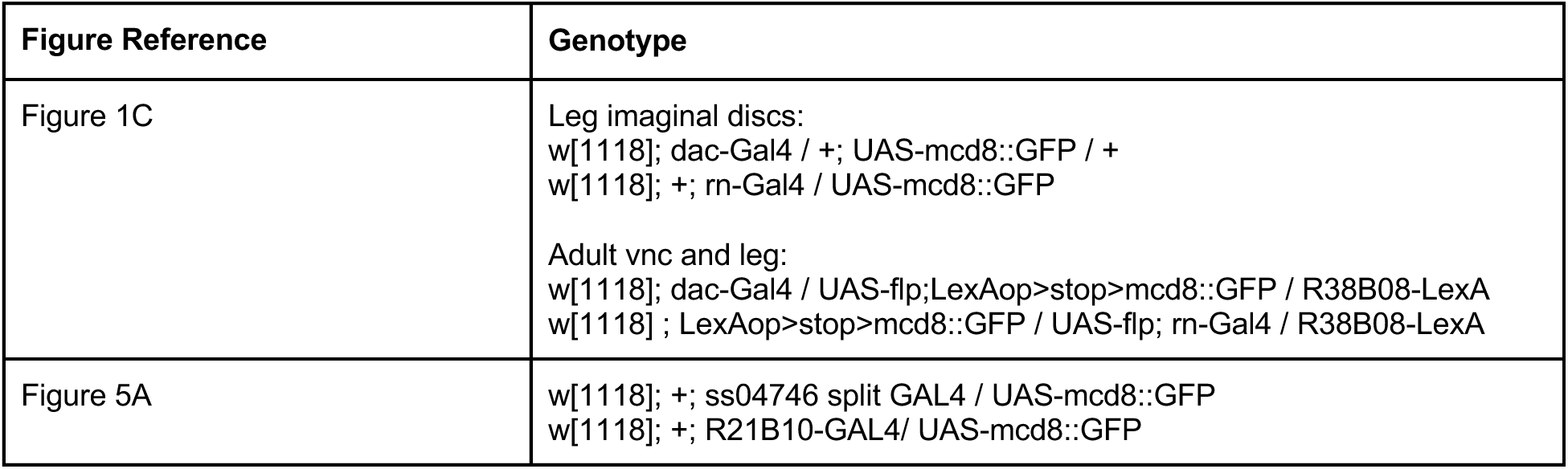
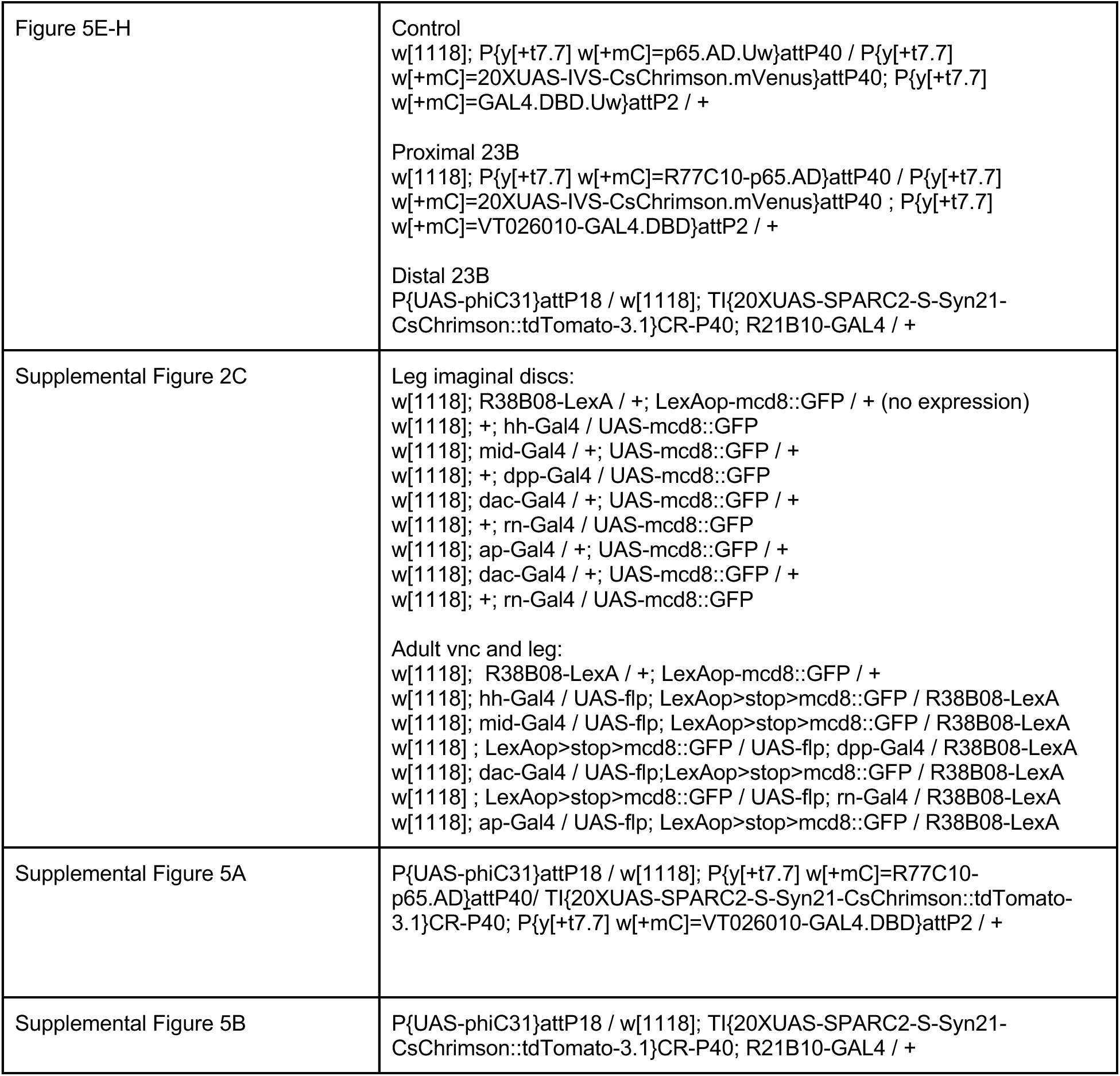

## Resource Availability

Code for analysis and figures can be found at: https://github.com/tuthill-lab/elabbady_bristles_2026

## Acknowledgements and Support

We thank members of the Tuthill Lab for technical assistance and continual feedback on the manuscript. We thank Elizabeth C. Marin and Haluk Lacin for their assistance with hemilineage identification for cells in FANC. We thank Casey Schneider-Mizell, Sven Dorkenwald, and Ben Pedigo for their thoughtful feedback on the manuscript, and Andy Seeds and Steffi Hampel for helpful discussions. We thank Igor Siwanowicz for sharing his blender model and his help understanding bristle distributions on the leg. We thank Kiet Tran for assistance proofreading neurons. L.E was supported by the Ruth L. Kirchstein Fellowship (F31NS134135) from the National Institute of Health. Other support was provided by the Allen Institute for Brain Science, the National Institutes of Health grants R01NS102333, R01NS128785, and U19NS104655, a Searle Scholar Award, a Klingenstein-Simons Fellowship, a Pew Biomedical Scholar Award, a McKnight Scholar Award, a Sloan Research Fellowship, the New York Stem Cell Foundation, and a UW Innovation Award to J.C.T. J.C.T is a New York Stem Cell Foundation – Robertson Investigator.

## Supplemental Figures

**Supplemental Figure 1:**
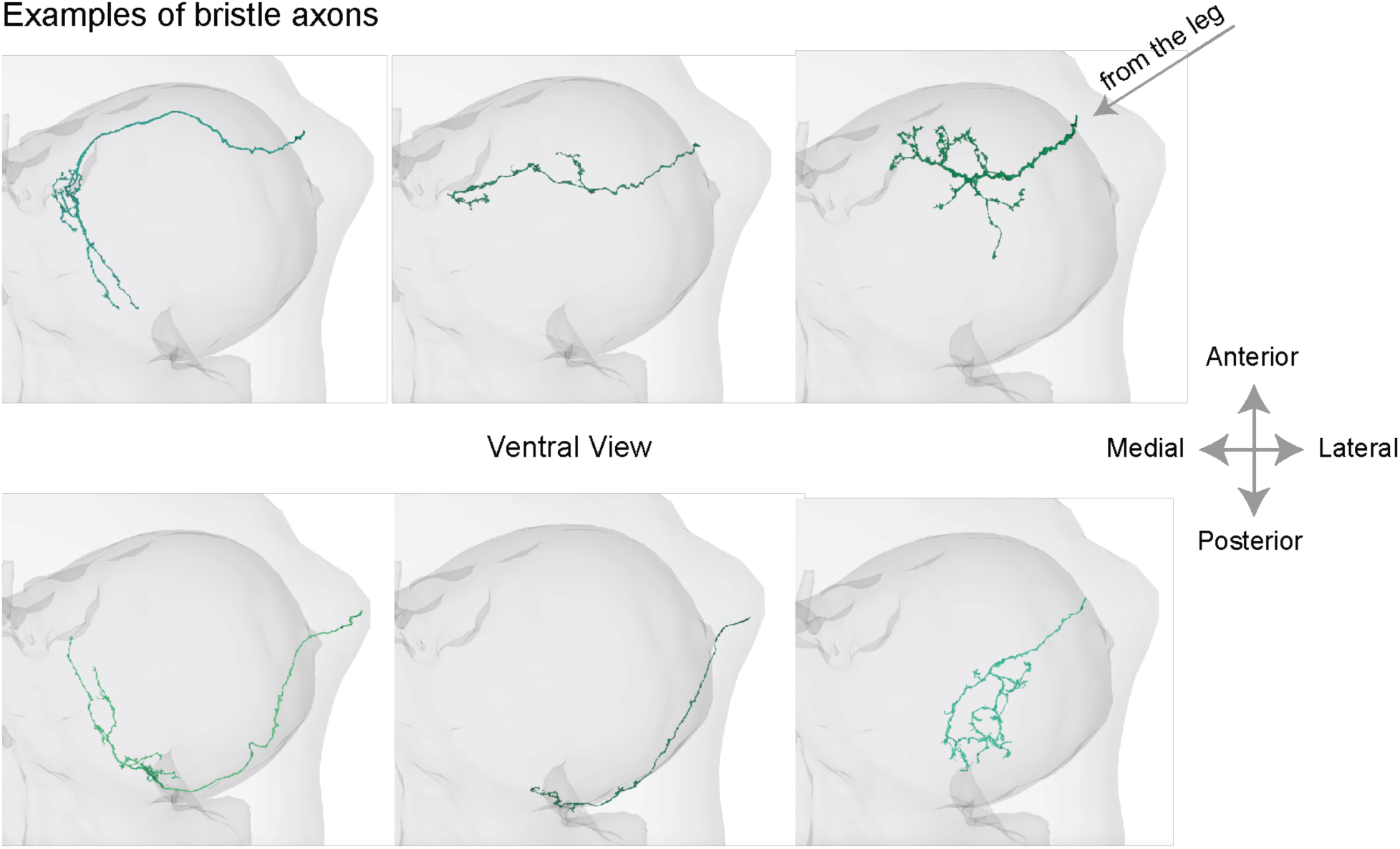
Bristle axons vary in morphology. Individual bristle axon morphologies. Three bristle axons that branch anteriorly (top row), and three that branch posteriorly (bottom row). Axons that cross the anterior to posterior border (left), axons that do not cross (middle), and axons that project closer to the center of the left leg neuromere (right).

**Supplemental Figure 2:**
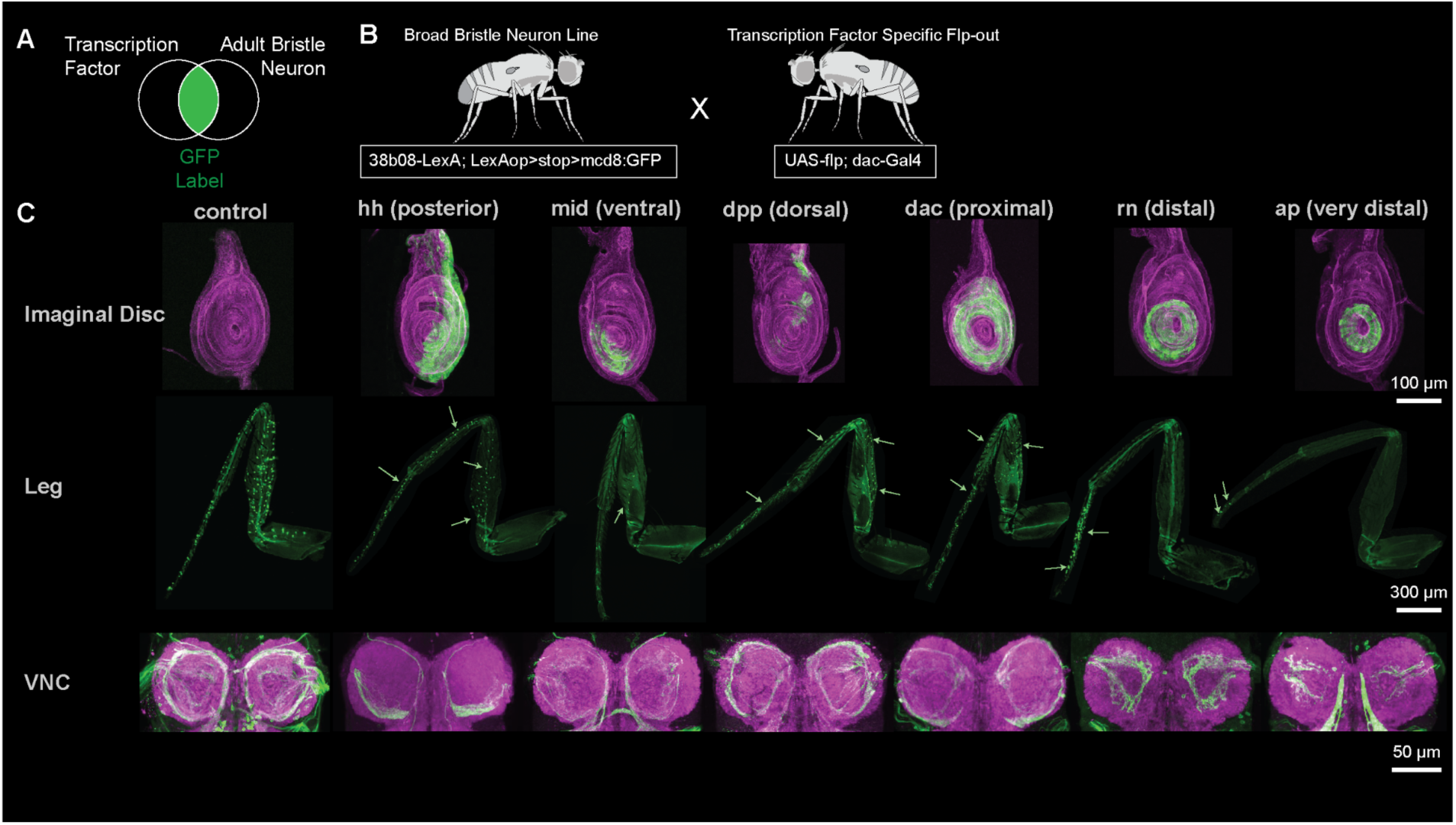
GFP expression of bristle neurons driven by coexpression of different transcription factors in the larval leg imaginal disc, leg, and VNC. **A)** For each line, only bristle cells that express a specific transcription factor will be labeled with GFP. **B)** Example genetic cross. **C)** Shown are maximum intensity projections of cells in the larval leg imaginal disc LexAop-mCD8::GFP(green) and an antibody against phalloidin (magenta). Bristle neurons in the leg and VNC were labeled with mcd8::GFP (green) and an antibody against the neuropil marker bruchpilot (magenta), green arrows indicate a sample of labeled bristle neurons. From left to right: all bristle neurons labeled by R38B08-LexA alone, bristle neurons that coexpressed *hedgehog* (hh), *midline* (mid), *decapentaplegic* (dpp), *dachshund* (dac), *rotund* (rn), and *apterous* (ap) during metamorphosis.

**Supplemental Figure 3:**
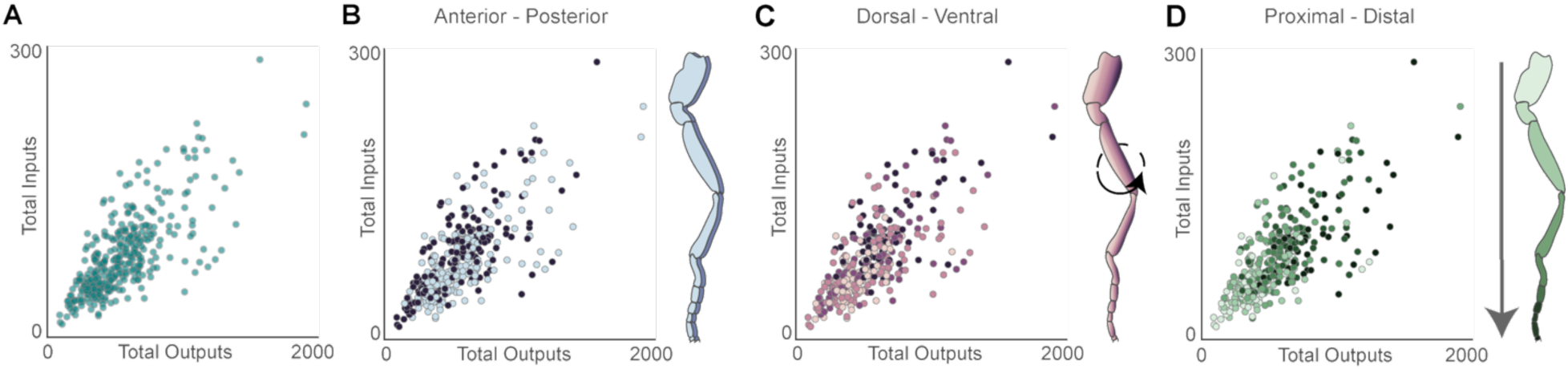
Synaptic input and output counts do not vary somatotopically. **A)** Number of input and output synapses for each reconstructed bristle axon (teal). Colored by the predicted spatial location on the leg along the **B)** anterior-posterior axis (r^2^=4.64e-05), **C)** dorsal-ventral axis (r^2^=0.05), **D)** proximal-distal axis (r^2^=0.30).

**Supplemental Figure 4:**
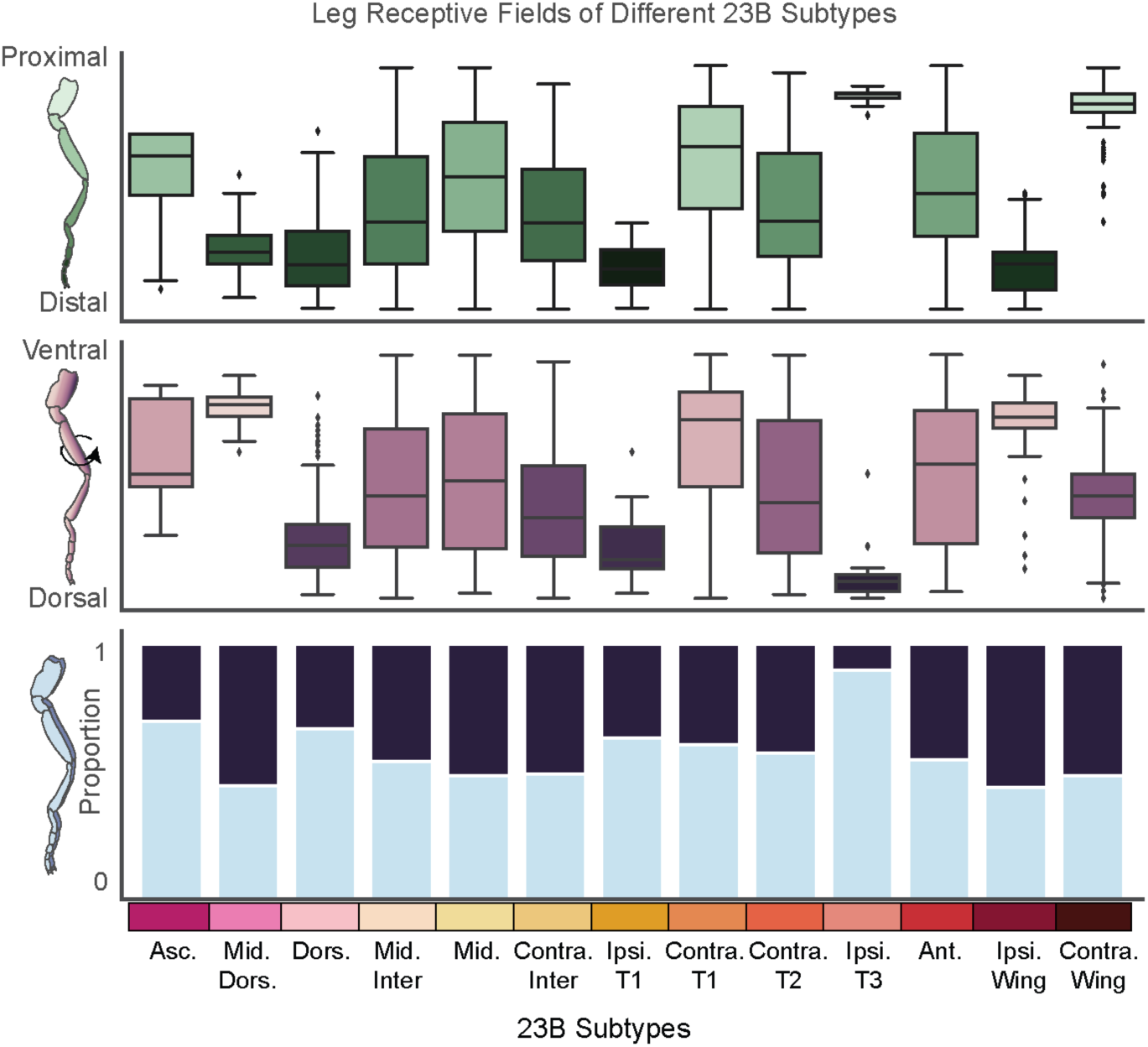
Leg receptive fields for different 23B subtypes. **A)** Receptive fields along the P/D axis (top), D/V axis (middle), A/P axis (bottom, anterior:light blue, posterior: dark blue) for different 23B subtypes. Individual points represent input synapses from bristle axons and the y axis represents where on the leg each presynaptic bristle axon originates. The order of 23B subtypes along the x axis is consistent across all three plots. For all box plots, center line, median; box limits, upper and lower quartiles; whiskers, 1.5x interquartile range.

**Supplemental Figure 5:**
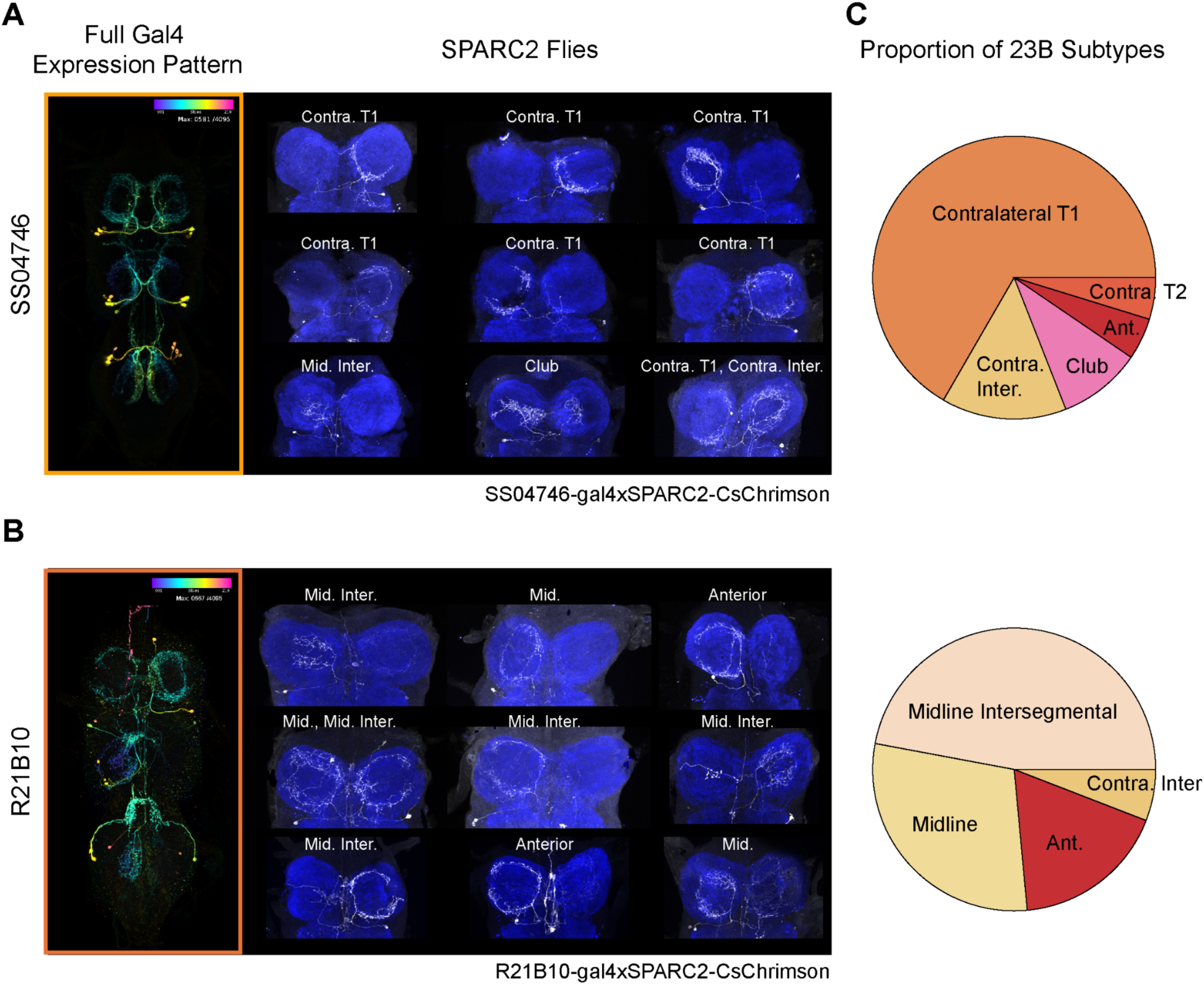
Experimental lines SS04746 and R21B10 label different 23B subtypes. **A)** Full Gal4 VNC expression of SS04746 from the Janelia library (left). Example VNCs from sparsified line SS04746-gal4xSPARC2-CsChrimson (right). **B)** Full Gal4 VNC expression of R21B10 from Janelia FlyLight^76^ (left). Example VNCs from sparsified line R21B10-gal4xSPARC2-CsChrimson (right). In all SPARC2 experiments 23B neurons in the VNC were labeled with mcd8::GFP (white) and an antibody against the neuropil marker bruchpilot (blue). Each neuron was classified by where in the VNC the axon projected to. **C)** Proportion of different 23B subtypes in SS04746 (n=21) and R21B10 (n=17).

**Supplemental Figure 6:**
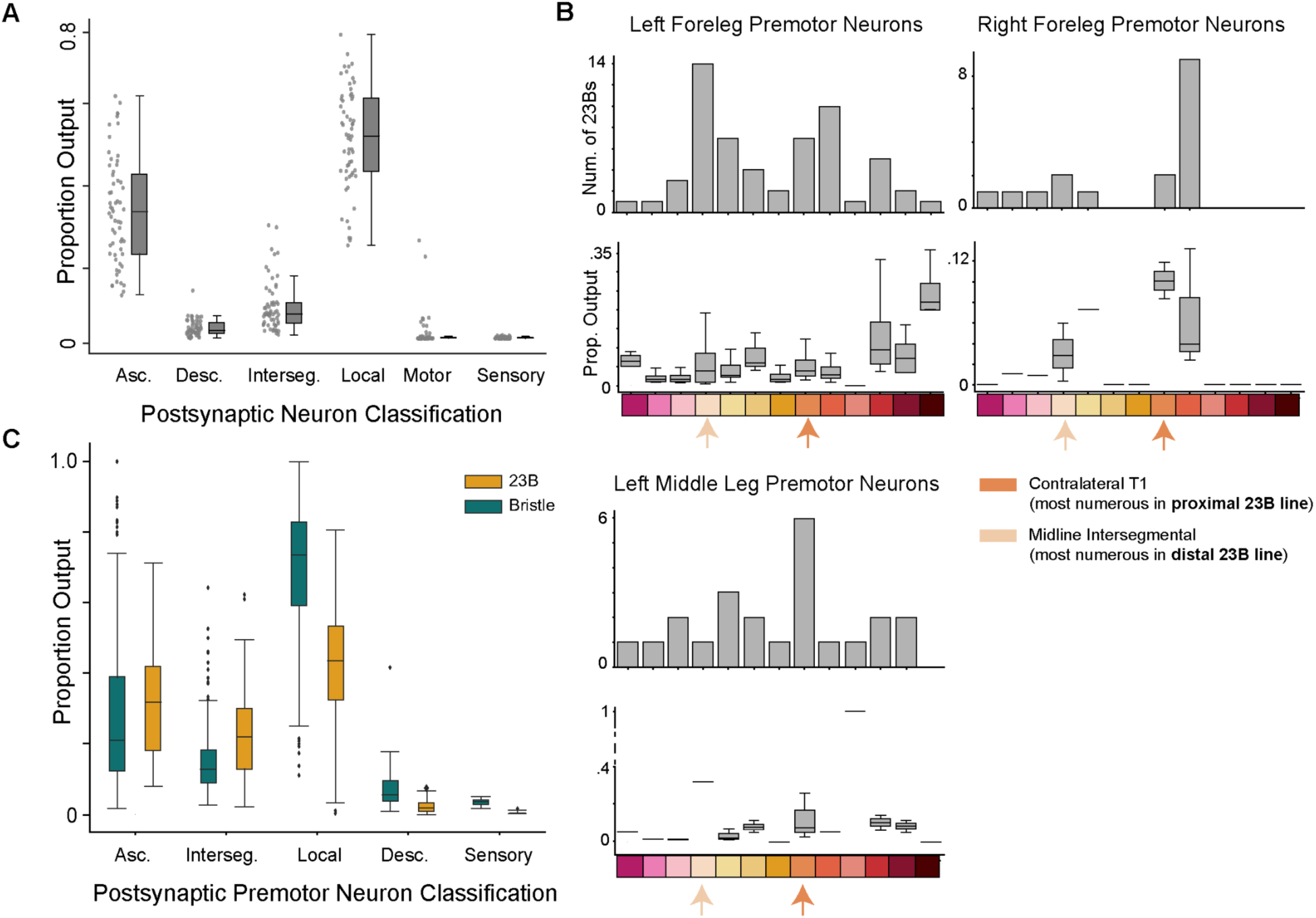
23B subtypes connectivity onto premotor neurons in T1L, T1R, and T2L. **A)** Proportion of 23B output connectivity onto different neuron classes in the VNC. **B)** 23B subtype connectivity onto premotor pools for the left front leg (T1L), right front leg (T1R), and the left middle leg (T2L). The bar graph represents the number of 23B neurons of each subtype that contact any premotor neurons within each leg neuropil. Boxplots represent the proportion of 23B output synapses onto premotor neurons within each leg neuropil. Color bars represent different 23B subtypes, from left to right: Ascending, Midline Dorsal, Dorsal, Midline Intersegmental, Midline, Contralateral Intersegmental, Ipsilateral T1, Contralateral T1, Contralateral T2, Ipsilateral T3, Anterior, Ipsilateral Wing, and Contralateral Wing. Arrows indicate the most prominent subtype in the *proximal-sensing* (SS04746) and *distal-sensing* (R21B10) grooming lines. **C)** Proportion of total premotor output onto different classification types; Asc: ascending, Desc: descending, Interseg: intersegmental. Color denotes the presynaptic partner either 23B (orange) or bristle neuron (teal). For all box plots, center line, median; box limits, upper and lower quartiles; whiskers, 1.5x interquartile range.

## Supplemental Videos

**Supplemental Video 1: *Proximal-sensing* 23B activation in headless flies.** Example trial for optogenetic activation of *proximal-sensing* 23B neurons (SS04746) expressing CsChrimson. Each trial was 20 seconds in duration, five seconds prestimulus, five seconds with the laser flickering on/off at 5Hz, and 10 seconds post stimulus.

**Supplemental Video 2: Distal-sensing 23B activation in headless flies.** Example trial for optogenetic activation of *distal-sensing* 23B neurons (R21B10) expressing CsChrimson. Each trial was 20 seconds in duration, five seconds prestimulus, five seconds with the laser flickering on/off at 5Hz, and 10 seconds post stimulus.

**Supplemental Video 3: Laser activation of empty-SpGal4 in headless flies.** Example trial for laser activation of empty-SpGal4 flies with no CsChrimson expression. Each trial was 20 seconds in duration, five seconds prestimulus, five seconds with the laser flickering on/off at 5Hz, and 10 seconds post stimulus.

